# The natural design for harvesting far-red light: the antenna increases both absorption and quantum efficiency of Photosystem II

**DOI:** 10.1101/2021.04.01.438080

**Authors:** Vincenzo Mascoli, Ahmad Farhan Bhatti, Luca Bersanini, Herbert van Amerongen, Roberta Croce

## Abstract

Cyanobacteria carry out photosynthetic light-energy conversion using phycobiliproteins for light harvesting and the chlorophyll-rich photosystems for photochemistry. While most cyanobacteria only absorb visible photons, some of them can acclimate to harvest far-red light (FRL, 700-800 nm) by integrating chlorophyll *f* and *d* in their photosystems and producing red-shifted allophycocyanin. Chlorophyll *f* insertion enables the photosystems to use FRL but slows down charge separation, reducing photosynthetic efficiency. Here we demonstrate with time-resolved fluorescence spectroscopy that charge separation in chlorophyll-*f*-containing Photosystem II becomes faster in the presence of red-shifted allophycocyanin antennas. This is different from all known photosynthetic systems, where additional light-harvesting complexes slow down charge separation. Based on the available structural information, we propose a model for the connectivity between the phycobiliproteins and Photosystem II that qualitatively accounts for our spectroscopic data. This unique design is probably important for these cyanobacteria to efficiently switch between visible and far-red light.

## Introduction

The light-dependent reactions of oxygenic photosynthesis are carried out by large assemblies of proteins, pigments, and electron carriers, known as Photosystem I (PSI) and Photosystem II (PSII) supercomplexes. These supercomplexes consist of an outer antenna performing light harvesting and a chlorophyll-rich photosystem core. The latter collects the excitations formed in the antenna and uses their energy to power charge separation through a pigment cluster in the centrally located reaction center (RC) complex^1^. While the photosystem cores, which are membrane-embedded, have a highly similar architecture in all oxygenic photoautotrophs, the antenna systems are more variable in both size and composition. This flexibility is required to cope with specific environmental conditions (such as the spectrum and/or intensity of light available)^2^.

While plants and most algae employ membrane-bound light-harvesting complexes^3^, cyanobacteria have water-soluble antennas, the phycobilisomes (PBSs)^4–6^. PBSs are relatively large complexes (typically several MDa) of phycobiliproteins containing covalently-bound chromophores named bilins, and associated linker proteins. Most cyanobacteria have a hemidiscoidal PBS consisting of a core composed of two, three, or five cylinders of allophycocyanin (APC) subunits, two of which are connected to the photosystem cores, and several rods of phycocyanin and, sometimes, phycoerythrin^7–9^. The downhill energy gradient between the peripheral rods and the APC core ensures directionality in the energy transfer towards the photosystems^10^. By binding up to several hundred pigments^11^, the PBSs enormously increase the absorption cross section of the photosystems and expand the photosynthetically active spectrum to regions (commonly between 500 and 650 nm) where the absorption by the chlorophylls (Chls) is weak. For most photosynthetic systems, however, a general consequence of using a larger antenna is a longer time required for the excitations to reach the RCs and power photochemistry, resulting in a lower photochemical efficiency^12^.

While most photoacclimation strategies pertain to the antenna, a group of cyanobacteria growing in deep-shaded environments can acclimate to harvest far-red light (FRL, 700-800 nm) by remodeling both their antenna and core photosystems^13^. Under visible light (such as white light, WL), these organisms produce only one type of Chl, Chl *a*, and absorb visible photons like “conventional” cyanobacteria. Under FRL, however, they become capable of harvesting less energetic quanta by synthesizing the red-shifted Chls *f* and *d*^13–15^. FRL-induced photoacclimation (FaRLiP) involves replacing the photosystems expressed in WL (WL-photosystems) with new photosystems containing FRL-specific paralog protein subunits. FRL-photosystems incorporate a small number of chlorophylls *f* and *d* allowing them to absorb up to 750 nm (FRL-PSII)^16–18^, or even 800 nm (FRL-PSI)^16,18–20^. Under FRL, new APC paralog subunits are also produced, which assemble into bicylindrical cores (FRL-BCs)^21,22^. While WL-PBSs absorb up to 670-680 nm, FRL-BCs contain a large number of red-shifted bilins absorbing above 700 nm^17,21,22^. The organization of phycobiliproteins in FaRLiP organisms is species-dependent. Some strains possess only FRL-BCs in FRL^22^, while in others the peripheral rods of phycocyanin and phycoerythrin are associated with FRL-BCs^23^. Finally, besides producing FRL-photosystems and FRL-BCs, some strains were shown to maintain variable amounts of WL-photosystems and/or WL-PBSs after weeks of acclimation to FRL^17,18,21^.

The use of red-shifted pigments represents a promising strategy for extending the photosynthetically active spectrum of other organisms, such as plants and algae. In order to achieve high biomass yields, however, the newly engineered photosynthetic units need to preserve high photochemical yields. In this respect, we have recently shown that the insertion of Chl *f* slows down photochemistry in both photosystems and significantly decreases the efficiency of FRL-PSII in comparison to WL-PSII^18^. This effect is ascribed to (i) a slower charge separation by the RCs due to a red-shifted primary electron donor and (ii) a less effective connectivity between the few FRL-absorbing Chls surrounded by a majority of Chls *a*. While the latter study investigated the two photosystems, little is known about the fate of the excitations harvested by their outer antenna, especially at physiological temperatures. This knowledge is particularly relevant because FRL-BCs absorb a substantial amount of FRL in the cells, implying that their performances have a considerable impact on the overall light-energy conversion. To shed more light on these aspects, we measured time-resolved fluorescence (TRF) of intact cells of the FaRLiP strain *Chroococcidiopsis thermalis* acclimated to FRL and compared these results with those from WL-acclimated cells. We performed experiments at two different excitation wavelengths to selectively excite the Chls or the bilins, thereby investigating the fate of the energy absorbed by either the photosystems or the phycobiliproteins. These data were successively used to quantify the efficiency of light conversion after exciting different units.

## Results

### Steady-state spectroscopy

In comparison with cells acclimated to WL (WL-cells), the cells acclimated to FRL (FRL-cells) display an additional absorption band above 700 nm (Figure 1A). This is caused by the integration of several Chls *f* in the FRL-photosystems and by the synthesis of red-shifted APC forming FRL-BCs^13^. The Chl content of FRL-cells is dominated by Chl *a* (∼95%). Chl *f* accounts for the remaining ∼5% (4.7±0.1%), whereas no Chl *d* could be detected. The extra absorption of FRL-cells in the blue-green region (λ < 530 nm) has also been observed in other FaRLiP strains acclimated to FRL and ascribed to an increased carotenoid content^15,19,24^.

**Figure 1.**
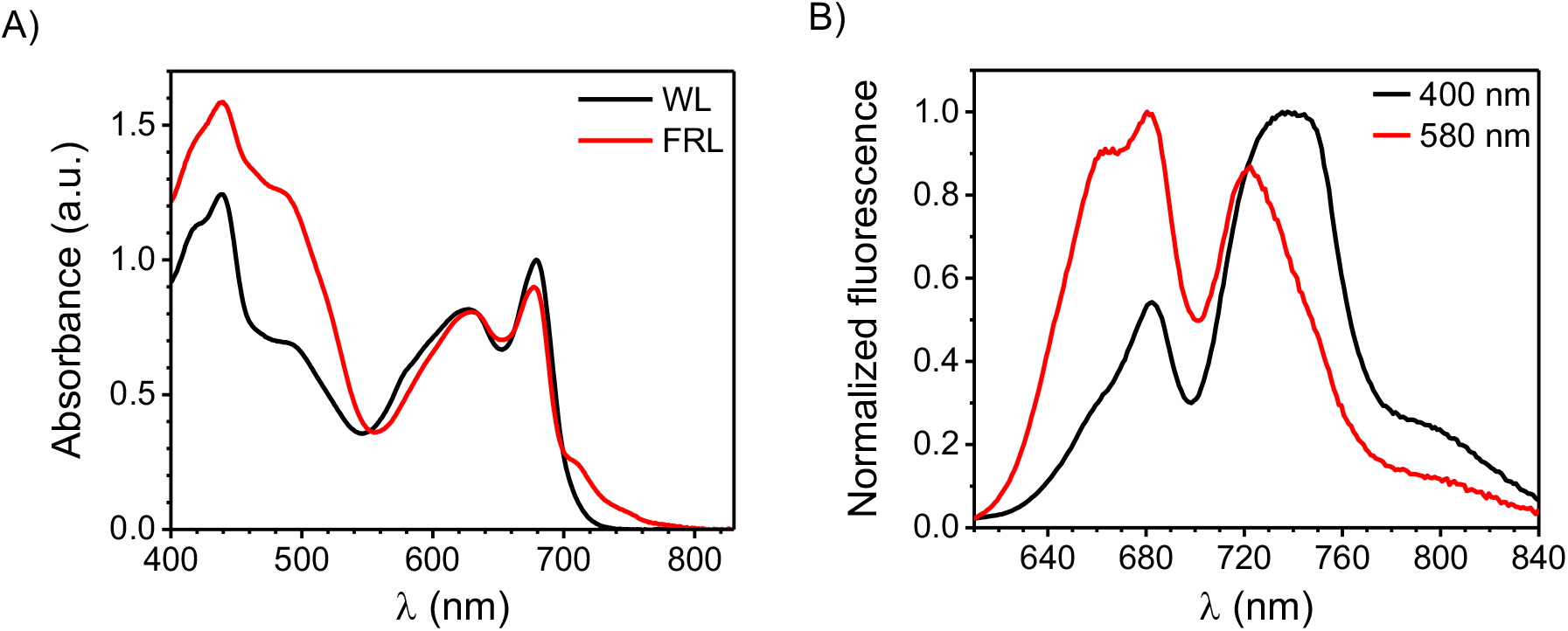
Steady-state spectra at room temperature of intact cells acclimated to WL or FRL. A) Absorption spectra of WL- and FRL-cells (normalized to the area in the region above 580 nm). B) Fluorescence spectra of FRL-cells excited at 400 and 580 nm. The corresponding spectra of WL-cells are displayed in Figure S1.

The fluorescence spectra of FRL-cells show two distinct bands upon excitation of both Chls (400 nm) and bilins (580 nm) (Figure 1B). The one at shorter wavelengths is due to WL-PSII and the WL-PBSs that are maintained in FRL, in line with previous findings on another FaRLiP strain^17^. The band above 700 nm is due to FRL-PSII and the FRL-BCs^18^, while the shoulder extending up to 800 nm stems from the extremely red-shifted Chls *f* typical for FRL-PSI^16,18,25^. Upon 580-nm excitation, which is more selective for the bilins, the far-red band is more blue-shifted than upon 400-nm excitation due to a larger contribution from FRL-BCs. The strong dependency of far-red fluorescence on the excitation wavelength indicates that FRL-BCs and FRL-photosystems do not fully equilibrate before excited-state decay. The 800-nm fluorescence is also substantially reduced when exciting at 580 nm, suggesting that the excited phycobiliproteins are less well connected to FRL-PSI than to other photosynthetic units.

### Trapping of Chl and bilin excitations by FRL-PSII RCs

To investigate the fate of different pigment excitations, the excited-state dynamics of intact cells were probed via TRF (Figures 2 and S2-9). Cells were excited both at low power, when most PSII RCs are in the open state, and at high power in the presence of 3-(3,4-dichlorophenyl)-1,1-dimethylurea (DCMU), when PSII RCs are in the closed state, to separate the spectroscopic contribution of PSII from that of PSI and estimate the photochemical yield of PSII. Experiments were performed at two different excitation wavelengths, 400 nm (more selective for the photosystems and yielding results similar to our previous work^18^) and 577 nm (more selective for the phycobiliproteins). Due to the heterogeneity of the photosynthetic apparatus of FRL-cells, both WL-PBSs and FRL-BCs are excited at 577 nm. Note that it is not possible to excite FRL-BCs selectively, as their absorption maxima (around 650 nm and 710 nm)^21,22^ overlap with the absorption bands of the Chls. However, an effective separation of the signatures from WL-PBSs and FRL-BCs is possible from the analysis of the spectrally-resolved TRF data. Below we only show the main findings for FRL-cells, whereas the study of WL-cells and the results from global analysis of all TRF datasets can be found in the Supplementary Materials (see Figures S3-8).

**Figure 2.**
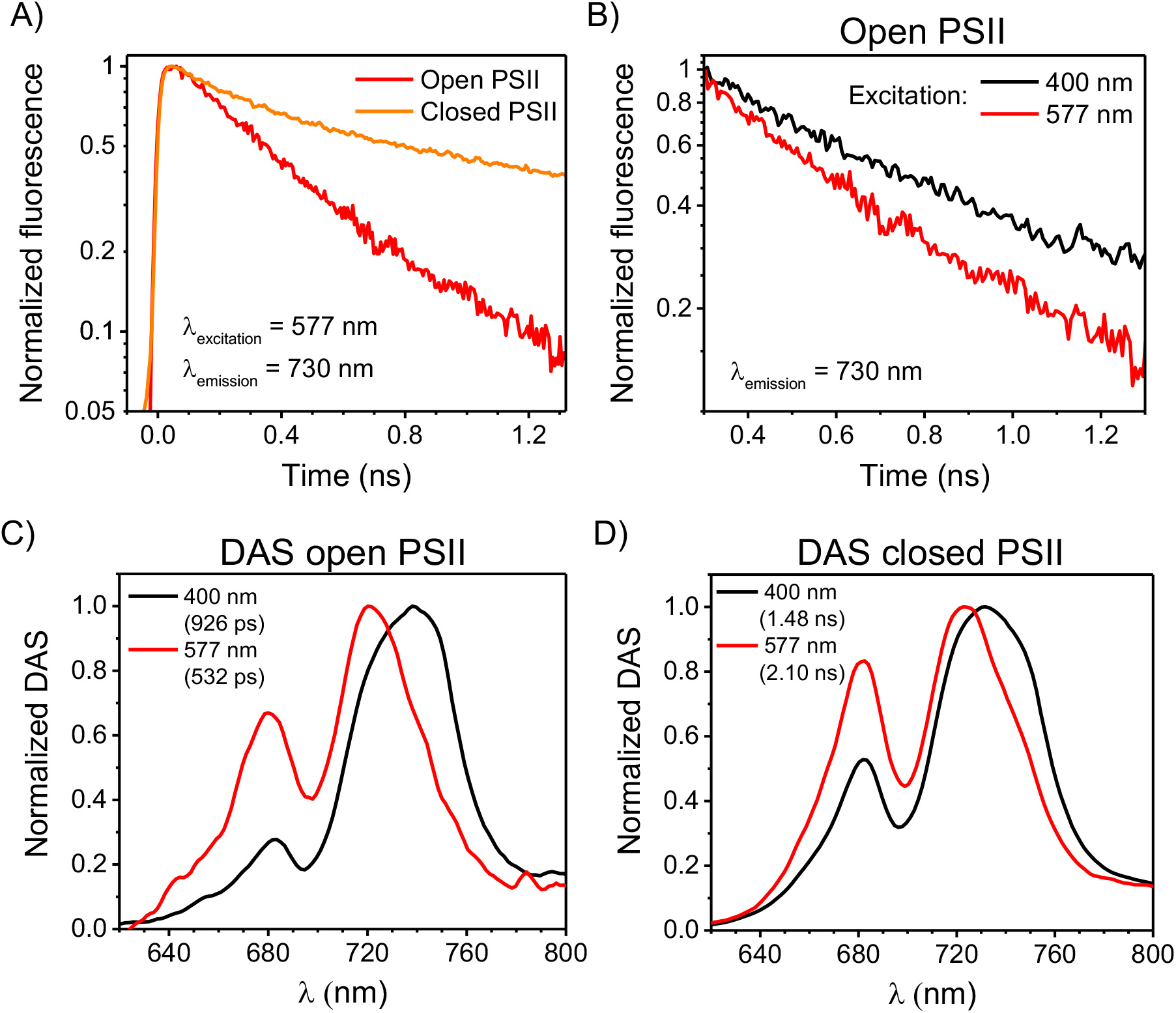
TRF of FRL-cells at room temperature. A) TRF traces (normalized to their maxima) detected at 730 nm (where both FRL-BCs and FRL-PSII emit) upon 577-nm excitation with mostly open and closed PSII RCs. The presented traces were obtained by integrating the TRF data between 725 and 735 nm and were binned to a time-step of 6 ps. B) TRF traces detected at 730 nm upon 400-nm and 577-nm excitation with mostly open PSII RCs. The traces are normalized to their value around 300 ps, and the first 300 ps of fluorescence kinetics are excluded for clarity. Please note that, in (A) and (B), the fluorescence intensity is shown on a logarithmic scale. C) Normalized long-lived DAS from TRF measurements with open PSII RCs after 400-nm and 577-nm excitation (corresponding to the green DAS in Figures S6A and S7A). D) Normalized long-lived DAS from TRF measurements with closed PSII RCs after 400-nm and 577-nm excitation (corresponding to the green DAS in Figure S6B and the magenta DAS in Figure S7B). The lifetimes of the DAS are indicated in parentheses in the legends. Note that, for the DAS in (C) and (D), the signal below 700 nm stems mostly from the WL-PSII and WL-PBSs that are maintained in the FRL-cells, whereas that above 700 nm is emitted by FRL-BCs and FRL-PSII.

For FRL-cells excited at 577 nm, the TRF traces detected at 730 nm (where both FRL-BCs and FRL-PSII emit) and normalized to their maxima become substantially longer-lived when PSII RCs close (Figure 2A). This indicates that FRL-BCs, which are more selectively excited at 577 nm, transfer their excitations mainly to FRL-PSII and are able to power photochemistry. The corresponding traces obtained upon 400-nm excitation, which is more selective for the photosystems, are shown in Figure S2. Note that, at earlier delays, the 400-nm-excited traces are dominated by the fast-decaying Chls *f* of FRL-PSI. As a result, the TRF traces of Figure 2A cannot be simply juxtaposed to those in Figure S2 to compare the trapping rates of Chl *f* and bilin excitations by PSII RCs. For a qualitative comparison, we therefore overlaid the 400-nm and 577-nm excited traces in open state normalized at a delay of 300 ps (Figure 2B). This delay is sufficiently longer than the timescale of energy transfer and trapping in FRL-PSI (< 200 ps, see also Figure S6)^18,26^, implying that the signal detected at 730 nm after 300 ps stems mainly from the FRL-BCs and FRL-PSII. When the 577-nm and 400-nm excited traces are normalized at this delay, the former trace is substantially shorter-lived. This implies that trapping of FRL-BCs excitations by FRL-PSII RCs is, on average, faster than trapping from the Chls *f*. The spectral signatures of FRL-BCs and FRL-PSII can be retrieved in the longest-lived decay-associated spectrum (DAS) obtained from global analysis of TRF data. The lifetime of the longest-lived DAS in open state, which can be used to estimate the time required to achieve charge separation, decreases from ∼930 ps upon 400-nm-excitation to ∼530 ps upon 577-nm excitation (Figure 2C; see Figures S6-7 for the full set of DAS). This confirms that energy trapping after red-shifted APC excitation is markedly faster than after Chl excitation. At the same time, the ∼930-ps DAS after preferential Chl excitation and the ∼530-ps DAS after bilin excitation do not overlap and their peaks are shifted by nearly 20 nm. This indicates that the former DAS stems mostly from Chls *f*, whereas the latter is enriched in red-shifted APC. A similar difference between the two excitations is observed for the longest-lived DAS when PSII RCs are closed (Figure 2D). These results imply that the excitations in the FRL-BCs and those in the FRL-PSII antenna do not fully equilibrate, even after 1-2 ns.

The association of FRL-BCs with FRL-PSII in FRL-acclimated *Chroococcidiopsis thermalis* appears to be different from that of PBSs and PSII core of previously studied cyanobacteria^27,28^ and WL-acclimated *Chroococcidiopsis thermalis*. In the latter case, WL-PSII RCs trap the excitations formed in WL-PBSs more slowly than those formed on the chlorophylls of the WL-PSII core. Furthermore, excitations of the bilins of WL-PBSs and of the Chls *a* of WL-PSII equilibrate after few hundred picoseconds (Figure S3).

### Efficiency of PSII photochemistry upon different pigment excitations

The TRF data presented above can be integrated to obtain steady-state fluorescence spectra in the open (F_o_) and closed (F_m_) states to calculate the variable fluorescence (F_v_ = F_m_ − F_o_) at various emission wavelengths (Figure 3). Variable fluorescence only stems from the pigments transferring to PSII RCs. The ratio F_v_/F_m_ in the spectral regions where PSII emission is maximal can be used to estimate the quantum yield of PSII photochemistry, Φ _PSII_ (see Figure S10 for details on the validity of this approximation).

**Figure 3.**
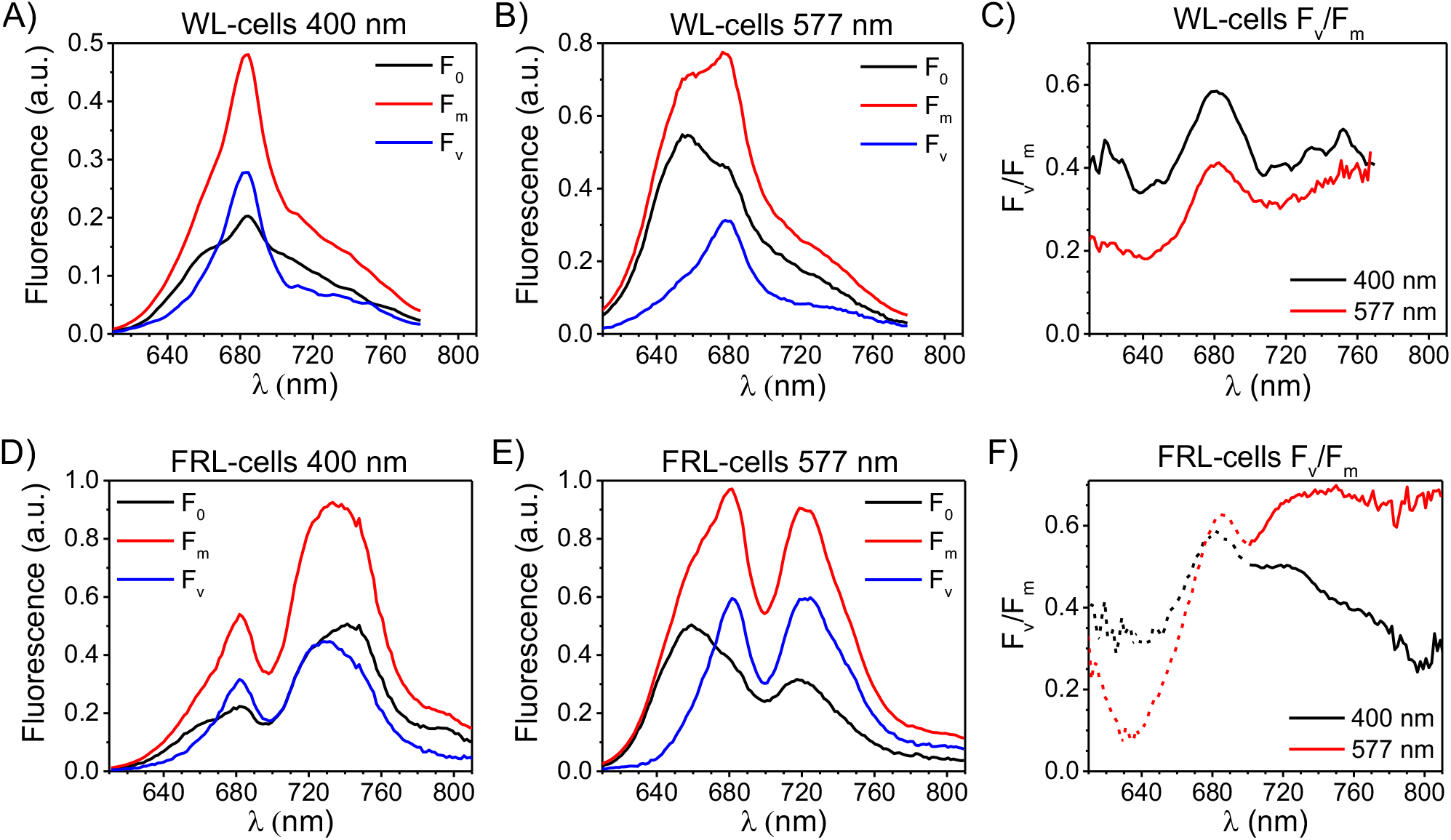
Variable fluorescence and F_v_/F_m_ of WL-and FRL-cells excited at different wavelengths at room temperature. A,B) Steady-state fluorescence spectra of WL-cells in open (F_o_) and closed (F_m_) state excited at 400 nm (A) and 577 nm (B) obtained from the DAS in Figures S4-5 and normalized to the area of the time-zero spectrum. Variable fluorescence spectra (F_v_) are calculated as the difference between F_m_ and F_o_. C) F_v_/F_m_ of WL-cells at different wavelengths calculated from the spectra in (A) and (B). D,E) Steady-state fluorescence spectra of FRL-cells in open (F_o_) and closed (F_m_) state excited at 400 nm (D) and 577 nm (E) obtained from the DAS in Figures S6-7. F) F_v_/F_m_ of FRL-cells at different wavelengths calculated from the spectra in (D) and (E). Note that the F_v_/F_m_ values around 680 nm upon 400-nm excitation in (F) are likely underestimated due to the presence of a small fraction of WL-PSII with closed RCs in the measurements at low powers (meaning that the integrated fluorescence in this region does not truly represent F_o_; see the DAS in Figure S6). As a result, these F_v_/F_m_ values cannot be directly compared to those with 577-nm excitation in the same region, which is why they are shown in dashed lines. On the other hand, FRL-PSII RCs are fully open in the measurements performed at low power with both 400-nm and 577-nm excitations (see Figure S11 for details). Consequently, the integrated fluorescence data of FRL-cells above 700 nm are representative of F_o_ and F_m_, and the corresponding F_v_/F_m_ values (continuous lines) can be used to compare the efficiencies of trapping by FRL-PSII RCs upon either Chl or bilin excitation.

At both excitation wavelengths (400 and 577 nm), the variable fluorescence spectrum of WL-cells peaks at ∼680 nm and stems from WL-PSII (Figures 3A and 3B), with a shoulder at shorter wavelengths due to the WL-PBS antenna. Upon 400-nm excitation, F_v_/F_m_ at 680 nm is nearly 0.6 (Figure 3C, black). This value of Φ_PSII_ is likely underestimated because both the excitation and emission wavelengths are not entirely selective for the Chls of WL-PSII. The maximal F_v_/F_m_ drops to ∼0.4 upon 577-nm excitation (Figure 3C, red), which is likely the result of two simultaneous effects: (i) not all WL-PBSs transfer energy to WL-PSII and (ii) trapping by WL-PSII after PBS excitation is slower than trapping after direct WL-PSII excitation due to the additional time required for the energy migration from the WL-PBS to WL-PSII. Consequently, WL-PBSs increase the antenna size of WL-PSII at the expense of some photochemical efficiency.

The variable fluorescence of FRL-cells shows two distinct peaks (Figures 3D and 3E): one around 680 nm, which we ascribe to WL-PSII and its WL-PBS antenna, and one above 700 nm due to FRL-PSII and its FRL-BC antenna. The F_v_ spectrum in the far-red region upon 400-nm excitation (Figure 3D, blue) is clearly blue-shifted with respect to both F_o_ and F_m_ (Figure 3D, black and red) and also relative to the spectrum of isolated FRL-PSII^18^. This indicates that trapping by FRL-PSII is more efficient at shorter wavelengths, where the contribution from FRL-BCs is larger. F_v_/F_m_ is therefore 0.50 at 720 nm (Figure 3F, black), it drops to about 0.44 at 740-750 nm, where fluorescence from the Chls *f* of FRL-PSII is maximal, and decreases even more at longer wavelengths, where FRL-PSI fluorescence dominates. Upon 577-nm excitation, as a consequence of the larger contribution from FRL-BCs, F_o_, F_m_ and F_v_ all peak around 720 nm in the far-red region (Figure 3E). In addition, Φ_PSII_ at λ > 700 nm is substantially higher than after 400-nm excitation, rising to a maximum of 0.68 (Figure 3F). All these results imply that FRL-PSII photochemistry after FRL-BCs excitation is more efficient than after direct Chl excitation, which is in contrast with what is normally observed for the supercomplexes of WL-PBS and WL-PSII of cyanobacteria^29^ (*cfr*. Figures 3C and 3F). The main cause of this somewhat counter-intuitive finding is that trapping by FRL-PSII RCs after excitation of FRL-BCs (requiring up to 500 ps) is remarkably faster than after Chl excitation (requiring up to 900 ps).

## Discussion

### Excitations formed in FRL-bicylindrical cores use a shortcut to reach the RCs of FRL-PSII

Our experimental results can be summarized as follows:

– In WL-cells, the PSII RCs trap excitations formed in the adjacent PBSs more slowly (and, therefore, less efficiently) than the excitations formed directly on the Chls. Furthermore, excitations in the WL-PBSs and WL-PSII equilibrate to a large extent within hundreds of picoseconds. These results are in line with what is commonly observed in cyanobacteria.
– Conversely, FRL-BCs can deliver excitations to the RC of FRL-PSII and power charge separation with a higher rate and efficiency than at least a fraction of the Chls *f* found in the FRL-PSII antenna (CP43 and CP47 paralogs). Furthermore, the excitations formed in the FRL-BCs do not equilibrate entirely with the Chls *f* of FRL-PSII, even after 2 ns. This behavior is different from most known photosynthetic systems/organisms and is, therefore, highly unexpected.

The lack of energy equilibration is usually ascribed to disrupted energetic connectivity between pigments. However, FRL-BCs display variable fluorescence and their excitations are trapped within 500 ps, implying that they are energetically well connected to the RCs of FRL-PSII. The only way to account for these seemingly contrasting results is to assume that, for structural reasons, the energy transfer route from FRL-BCs to the RC of FRL-PSII is faster than that connecting some of the Chls *f* in the antenna and the RC itself. This also explains why the excitations formed in the FRL-BCs can reach the RC without equilibrating with all the Chls *f* in the FRL-PSII antenna. The shortcut connecting the FRL-BCs to the FRL-PSII RCs is likely to involve an intermediate Chl in the antenna (CP43/47 paralogs) acting as a bridge. Indeed, structural data on canonical PBSs have shown that the pigments in the APC core are too far from those of the PSII RC to allow direct energy transfer between them^8,9,30^. To allow for faster trapping of bilin excitations, this bridging Chl in the antenna needs to be better connected to the RC than most other red-shifted Chls and, to avoid energetic barriers when receiving excitations from FRL-BCs, it is likely to be red-shifted as well.

This peculiar kinetics is the result of the small number of red-shifted pigments surrounded by a majority of Chls *a*. In light of this architecture, the connectivity between the FRL-PSII antenna and the RC can only count on the few Chls *f* in CP43/47, which rapidly harvest all excitations from the surrounding Chls *a*^18^, and the single red-shifted pigment in the RC, most likely the primary donor, Chl_D1_^16^. The couplings between these Chls are relatively weak and, as a result, the overall energy migration towards the RC is slowed down relative to WL-PSII^18^. At the same time, the fluorescence decay of purified FRL-PSII in open state is, however, multi-exponential^18^: this can be interpreted in terms of the RCs trapping excitations from different red-shifted Chls in the antenna on different timescales, from 100-200 ps to about 1 ns. Chl_D1_ is spatially closer to CP43^31^ and should therefore trap excitations from its Chls *f* more rapidly than from CP47 (if any). Indeed, based on the calculated couplings, the only Chl able to transfer excitations to Chl_D1_ in less than 100 ps (if occupied by a red-shifted Chl) is Chl505 of CP43 (Chl nomenclature based on the structure with PDB code 3WU2^31^), whereas five more Chls of CP43 and two of CP47 could transfer in 200-400 ps (if Chls *f*, see Figure 4). The lifetime heterogeneity of FRL-PSII could therefore be explained by a faster trapping of the excitations from the Chls *f* of CP43 and a slower trapping from the Chls *f* of CP47 by the RC. By contrast, the migration time in canonical PSII is in the order of 50 ps or less, with no large differences between CP43 and CP47^32,33^. In light of this hypothesis, Chl505 of CP43 represents an ideal candidate for bridging the FRL-BCs to FRL-PSII RCs. Not only is Chl505 the most strongly coupled pigment to Chl_D1_, it is also located at the stromal side of the complex, where the PBS basal core cylinder attaches to PSII. Furthermore, structural data/models of supercomplexes of PBS and (Chl *a*-only) PSII, indicate that the terminal emitters of the PBS, located in the ApcE subunit, are spatially closer to the Chls of CP43 than those of CP47^8,34^. Finally, the weak connectivity between the Chls of CP43 and those of CP47 could account for the dependency of the fluorescence spectra on the excitation wavelength. Indeed, the excitations formed on FRL-BCs would only equilibrate with the Chls of CP43 without reaching those of CP47, and vice versa. Notably, a recent work has left space for another RC chlorophyll, P_D2_, to function as the red-shifted primary donor instead of Chl_D1_^35^, though this possibility seems less likely due to mechanistic and structural considerations. However, even when the primary donor is assumed to be located on P_D2_, Chl 505 of CP43 remains the most probable candidate for connecting the FRL-BCs to FRL-PSII RCs (see Table S1 for details).

**Figure 4.**
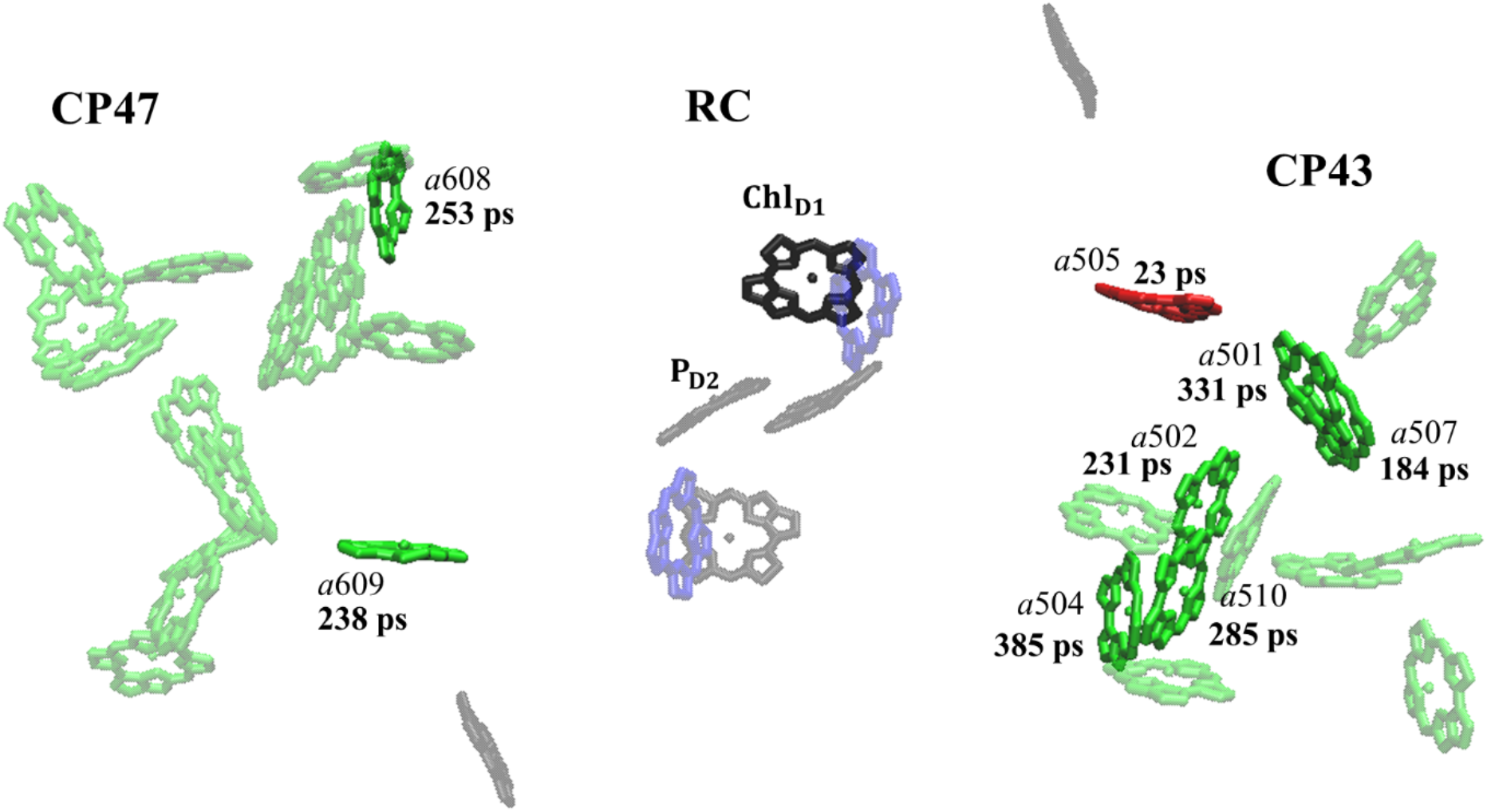
Hopping times between potential binding sites for red-shifted Chls in the antenna and the primary donor Chl_D1_. The Chl binding sites are taken from the PSII structure of *T. vulcanus* by Umena et al.^31^. The hopping times (inverse of energy transfer rates) are taken from Table S1 and are calculated in the case where the energy donor in the antenna is a Chl *f* emitting at 740 nm (based on the average emission wavelength of FRL-PSII) and the acceptor (Chl_D1_) is a Chl *f* absorbing at 727 nm (based on Nurnberg et al.)^16^. The RC Chls are shown in transparent black, and those in the antenna (CP43/47) in transparent green. Chl_D1_ is shown in solid black, while the antenna Chls with a hopping time below 500 ps are in solid green. Chl505, which exhibits the fastest calculated energy transfer towards Chl_D1_, is colored in red. The two pheophytin molecules in the RC are displayed in transparent blue.

As a result of the unusual architecture and energetics, FRL-BCs increase both the antenna size and the photochemical yield of FRL-PSII RCs simultaneously, which represents a unique case in photosynthetic systems. Based on the absorption spectra of the isolated complexes, it can be estimated that FRL-BCs contain about twice as many FRL-absorbing pigments (∼20 bilins; see Figure S12 for further details) as the FRL-PSII dimer (which binds ∼10 Chls *f/d*)^16–18^. As a result, they increase the antenna size of FRL-PSII under FRL by a factor of 3. At the same time, FRL-BC excitations are processed 1.5 times more efficiently than Chl excitations (on average, see previous section), implying that FRL-BCs could increase the overall performance of FRL-PSII RCs by a factor of 4 in FRL. This finding can also shed light on some other peculiar features of FaRLiP. Indeed, while most photoacclimation strategies developed by photosynthetic organisms are restricted to the antenna, FaRLiP also remodels the photosystem cores, which change their protein and pigment composition^13^. The insertion of red-shifted pigments is not particularly advantageous in terms PSII efficiency^18^, but might be needed to ensure energetic connectivity between the FRL-BC antenna, which contains the majority of FRL-absorbing pigments, and the RC. Indeed, a Chl *a*-only PSII core would hardly trap excitations from such a red-shifted antenna. Notably, FaRLiP is also substantially different from the strategy adopted by the only other known cyanobacterium able to grow in FRL, *Acaryochloris marina*^36^. This species also uses the red-shifted Chl *d* but, unlike FaRLiP strains, where Chl *a* remains the most abundant pigment also in FRL, *Acaryochloris* photosystems bind mostly Chl *d*^36–38^. The almost exclusive presence of Chl *d* results in a better energetic connectivity in the PSII of *Acaryochloris*, which shows therefore higher photochemical yields than FRL-PSII of FaRLiP strains^18^. At the same time, however, the phycobiliproteins produced by *Acaryochloris* only absorb visible light^39^, whereas FaRLiP strains harvest a substantial amount of FRL via red-shifted APC. In this view, the positioning of the long-wavelength Chls might be optimized for FRL-PSII to work in combination with FRL-BCs rather than on its own. In addition, the observation that the antenna compensates for the relatively low efficiency of FRL-PSII alone might also partially explain why the latter does not accumulate in FaRLiP organisms depleted in FRL-absorbing APC^40^.

Why do FaRLiP strains and *Acaryochloris* adopt such different strategies to convert low-energy photons? One possible explanation is that FaRLiP is an acclimative process allowing cyanobacteria to harvest FRL only when needed, whereas *Acaryochloris* employs Chl *d* permanently. The necessity to switch back to the more efficient Chl *a*-only photosystems when visible light becomes available might explain why FaRLiP cyanobacteria use Chl *a* as their major Chl also in FRL. As a result, the insertion of only few red-shifted Chls in the PSII core while most far-red photons are collected by the outer antenna would make FaRLiP cyanobacteria more flexible than *Acaryochloris* towards changes in the light spectrum. This hypothesis is also in line with the fact that FaRLiP has been adopted by many distantly-related species populating a variety of terrestrial and marine environments, whereas the strategy adopted by *Acaryochloris* remains unique^15,41^. Finally, the use of red-shifted Chls allows harvesting less energetic photons but also introduces additional challenges for the photosystems (such as a lower efficiency, and/or a higher probability of charge recombination)^16,18^, which is why it is restricted to environments enriched in FRL.

## Materials and Methods

### Cell cultures

The strain *Chroococcidiopsis thermalis* PCC 7203 was obtained from the Pasteur Culture Collection (Institut Pasteur, Paris, France) and grown at 30 °C in BG11^42^ medium with addition of 20 mM HEPES-NaOH (pH = 8.0). Cells were grown under white light (WL) of 30 µmol photons m^-2^ s^-1^ starting from OD_750_ = 0.1 and collected for experiments at OD_750_ = 0.8-1.0. Cells were also grown under far-red light (FRL, 738 nm; Jazz) of 45 µmol photons m^-2^ s^-1^ for 2.5 to 3 weeks prior to experiments, starting from OD_750_ = 0.3-0.4, with the medium being refreshed every week keeping an OD_750_ = 1.0-1.5. The cultures were grown in Erlenmeyer flasks shaken at 100 rpm. All measurements were performed on intact cells in their logarithmic growth phase.

### Chlorophyll content determination

Pigments from FRL-cells were extracted in pure methanol and their concentrations in the extract were determined by absorbance measurements. To improve the extraction efficiency of methanol, glass beads (about 30% v/v) were added and the mixture was vigorously shaken for 30 min in darkness. The absorption spectra of the Chls in the Q_y_ region were fitted using the available spectra and extinction coefficients of the pure pigments in methanol^43^. Average relative concentrations (and standard deviations) of the Chls were obtained from 4 measurements on 2 distinct cell batches. Note that the presence of WL-photosystems (particularly WL-PSII) in FRL-cells has an influence on the overall Chl content.

### Steady-state spectroscopy

Room-temperature (RT) absorption spectra of cells were acquired on a Varian Cary 4000 UV–Vis-spectrophotometer (Agilent technologies) equipped with an integrating diffuse reflectance sphere (DRA-CA-50, Labsphere) to correct for light scattering. RT fluorescence spectra were acquired at an OD < 0.05 cm^-1^ at Q_y_ maximum on a HORIBA Jobin-Yvon FluoroLog-3 spectrofluorometer. Absorption and fluorescence spectra were measured on at least three biological replicates for both FRL- and WL-acclimated cells, yielding similar results.

### Time-resolved fluorescence

Time-resolved fluorescence (TRF) measurements with 400-nm excitation were performed with a synchro scan streak camera setup as in ref. ^18^. TRF measurements with 577-nm excitation were recorded with a slightly different synchro scan streak camera setup^44,45^. In brief, the laser repetition rate was 3.8 MHz and the time window 2.0 ns. 577-nm excitation measurements with nearly all PSII RCs open were performed with an excitation power of 0.2 µW (with a sample volume of 2 mL), while those with closed PSII RCs were performed at 2 µW, after addition of 3-(3,4-dichlorophenyl)-1,1-dimethylurea (DCMU) and pre-illumination with white light for one minute (with a sample volume of 1 mL). The experiments were performed at RT in a magnetically stirred 1 cm × 1 cm cuvette with a sample OD of about 0.5 cm^-1^ at Q_y_ maximum (excitation and detection from the sample at the edge of the cuvette, thereby avoiding self-absorption) and a measuring time from 15 minutes to 1 hour of CCD exposure (no sample degradation was observed during the measuring time). The averaged images were corrected for background and shading, and then sliced into traces of ∼1.5-nm width prior to analysis. 400-nm excitation experiments consisted of three (two) biological replicas for FRL- (WL-) adapted cells, while 577-nm experiments consisted of two (one) biological replicas for FRL- (WL-) adapted cells. The biological replicas yielded very similar results. All TRF experiments were performed at RT.

### Data analysis

Fluorescence time traces were globally analyzed with Glotaran and the TIMP package for R^46^ using a number of parallel kinetic components. The total dataset can be described by the fitting function *f*(*t, λ*):

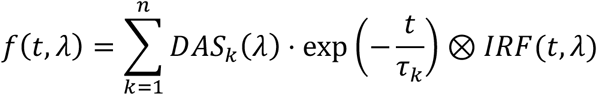

where each decay-associated spectrum (*DAS*_*k*_) is the amplitude factor associated with a decay component *k* having a decay lifetime *τ*_*k*_. The instrument response function *IRF*(*t, λ*) was estimated from the fitting (FWHM ∼ 20 ps) using a Gaussian profile. The fitting also accounts for time-zero dispersion. In some cases, a double Gaussian was required, consisting of a ∼20 ps FWHM (90% of IRF area) on top of a Gaussian of ∼100 ps FWHM (10% of IRF area). For each experiment, the time-zero spectrum was obtained by summing all DAS. The steady-state fluorescence spectra were reconstructed by integrating the TRF data as:

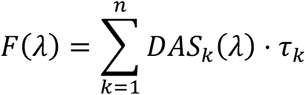

In order to compare results from different measurements, the DAS and reconstructed steady-state fluorescence spectra were normalized to the area of the time-zero spectrum (corresponding to the same amount of initial excitation) in each dataset. Overlays of the raw and fitted kinetic traces can be found in Figure S9.

## Acknowledgments

This project was supported by the European Union’s Horizon 2020 research and innovation program under the Marie Skłodowska-Curie grant agreement no. 675006 (to HvA and RC) and by the Netherlands Organization for Scientific Research (NWO) via a Top grant to RC. LB was supported by an EMBO long-term fellowship (EMBO ALTF 292-2017).

## Data and materials availability

All data needed to evaluate the conclusions in this paper are present in the paper and/or Supplementary Materials. All original data are available from the authors upon reasonable request.

## References

1. Croce, R. & van Amerongen, H. Light harvesting in oxygenic photosynthesis: structural biology meets spectroscopy. Science 369,933 (2020).

2. Croce, R. & van Amerongen, H. Natural strategies for photosynthetic light harvesting. Nat. Chem. Biol. 10,492–501 (2014).

3. Pan, X., Cao, P., Su, X., Liu, Z. & Li, M. Structural analysis and comparison of light-harvesting complexes I and II. Biochim. Biophys. Acta - Bioenerg. 1861,148038 (2020).

4. Bryant, D. A. & Canniffe, D.P. How nature designs light-harvesting antenna systems: design principles and functional realization in chlorophototrophic prokaryotes. J. Phys. B At. Mol. Opt. Phys. 51,33001 (2018).

5. Adir, N. Light harvesting in cyanobacteria: the phycobilisomes. in Light Harvesting in Photosynthesis 77–94 (CRC Press, 2018).

6. Adir, N., Bar-Zvi, S. & Harris, D. The amazing phycobilisome. Biochim. Biophys. Acta - Bioenerg. 1861,148047 (2020).

7. Arteni, A. A., Ajlani, G. & Boekema, E.J. Structural organisation of phycobilisomes from Synechocystis sp. strain PCC6803 and their interaction with the membrane. Biochim. Biophys. Acta - Bioenerg. 1787,272–279 (2009).

8. Chang, L. et al. Structural organization of an intact phycobilisome and its association with photosystem II. Cell Res. 25,726–737 (2015).

9. Liu, H. et al. Phycobilisomes supply excitations to both photosystems in a megacomplex in cyanobacteria. Science 342,1104–1107 (2013).

10. van Grondelle, R., Dekker, J. P., Gillbro, T. & Sundström, V. Energy transfer and trapping in photosynthesis. Biochim. Biophys. Acta 1187,1–65 (1994).

11. Ma, J. et al. Structural basis of energy transfer in Porphyridium purpureum phycobilisome. Nature 579,146–151 (2020).

12. van Amerongen, H., Valkunas, L. & van Grondelle, R. Photosynthetic excitons. (World Scientific Publishing, 2000).

13. Gan, F. et al. Extensive remodeling of a cyanobacterial photosynthetic apparatus in far-red light. Science 345,1312–1317 (2014).

14. Chen, M., Li, Y., Birch, D. & Willows, R.D. A cyanobacterium that contains chlorophyll f - a red-absorbing photopigment. FEBS Lett. 586,3249–3254 (2012).

15. Gan, F., Shen, G. & Bryant, D.A. Occurrence of far-red light photoacclimation (FaRLiP) in diverse cyanobacteria. Life 5,4–24 (2015).

16. Nürnberg, D. J. et al. Photochemistry beyond the red limit in chlorophyll f-containing photosystems. Science 360,1210–1213 (2018).

17. Ho, M.-Y. et al. Extensive remodeling of the photosynthetic apparatus alters energy transfer among photosynthetic complexes when cyanobacteria acclimate to far-red light. Biochim. Biophys. Acta - Bioenerg. 1861,148064 (2020).

18. Mascoli, V., Bersanini, L. & Croce, R. Far-red absorption and light-use efficiency trade-offs in chlorophyll f photosynthesis. Nat. Plants 6,1044–1053 (2020).

19. Li, Y., Vella, N. & Chen, M. Characterization of isolated photosystem I from Halomicronema hongdechloris, a chlorophyll f-producing cyanobacterium. Photosynthetica 56,306–315 (2018).

20. Kurashov, V. et al. Energy transfer from chlorophyll f to the trapping center in naturally occurring and engineered photosystem I complexes. Photosynth. Res. 141,151–163 (2019).

21. Ho, M.-Y., Gan, F., Shen, G. & Bryant, D.A. Far-red light photoacclimation (FaRLiP) in Synechococcus sp. PCC 7335. II. Characterization of phycobiliproteins produced during acclimation to far-red light. Photosynth. Res. 131,187–202 (2017).

22. Li, Y. et al. Characterization of red-shifted phycobilisomes isolated from the chlorophyll f-containing cyanobacterium Halomicronema hongdechloris. Biochim. Biophys. Acta - Bioenerg. 1857,107–114 (2016).

23. Soulier, N., Laremore, T. N. & Bryant, D.A. Characterization of cyanobacterial allophycocyanins absorbing far-red light. Photosynth. Res. (2020). doi:10.1007/s11120-020-00775-2

24. Schmitt, F. J. et al. Photosynthesis supported by a chlorophyll f-dependent, entropy-driven uphill energy transfer in Halomicronema hongdechloris cells adapted to far-red light. Photosynth. Res. 139,185–201 (2019).

25. Tros, M. et al. Breaking the red limit: efficient trapping of long-wavelength excitations in chlorophyll-f-containing Photosystem I. Chem 7,1–19 (2021).

26. Kaucikas, M., Nürnberg, D., Dorlhiac, G., Rutherford, A. W. & van Thor, J.J. Femtosecond visible transient absorption spectroscopy of chlorophyll f-containing photosystem I. Biophys. J. 112,234–249 (2017).

27. Tian, L. et al. Site, rate, and mechanism of photoprotective quenching in cyanobacteria. J. Am. Chem. Soc. 133,18304–18311 (2011).

28. Akhtar, P. et al. Time-resolved fluorescence study of excitation energy transfer in the cyanobacterium Anabaena PCC 7120. Photosynth. Res. 144,247–259 (2020).

29. Remelli, W. & Santabarbara, S. Excitation and emission wavelength dependence of fluorescence spectra in whole cells of the cyanobacterium Synechocystis sp. PPC6803: influence on the estimation of photosystem II maximal quantum efficiency. Biochim. Biophys. Acta - Bioenerg. 1859,1207–1222 (2018).

30. Krasilnikov, P. M., Zlenko, D. V. & Stadnichuk, I.N. Rates and pathways of energy migration from the phycobilisome to the photosystem II and to the orange carotenoid protein in cyanobacteria. FEBS Lett. 594,1145–1154 (2020).

31. Umena, Y., Kawakami, K., Shen, J. R. & Kamiya, N. Crystal structure of oxygen-evolving photosystem II at a resolution of 1.9 Å. Nature 473,55–60 (2011).

32. Raszewski, G. & Renger, T. Light harvesting in photosystem II core complexes is limited by the transfer to the trap: can the core complex turn into a photoprotective mode?J. Am. Chem. Soc. 130,4431–4446 (2008).

33. van der Weij-de Wit, C. D., Dekker, J. P., van Grondelle, R. & van Stokkum, I. H. M. Charge separation is virtually irreversible in photosystem II core complexes with oxidized primary quinone acceptor. J. Phys. Chem. A 115,3947–3956 (2011).

34. Zlenko, D. V., Krasilnikov, P. M. & Stadnichuk, I.N. Structural modeling of the phycobilisome core and its association with the photosystems. Photosynth. Res. 130,347– 356 (2016).

35. Judd, M. et al. The primary donor of far-red photosystem II: ChlD1 or PD2? Biochim. Biophys. Acta - Bioenerg. 1861,148248 (2020).

36. Miyashita, H. et al. Chlorophyll d as a major pigment. Nature 383,402 (1996).

37. Tomo, T. et al. Characterization of highly purified photosystem I complexes from the chlorophyll d-dominated cyanobacterium Acaryochloris marina MBIC 11017. J. Biol. Chem. 283,18198–18209 (2008).

38. Allakhverdiev, S. I. et al. Redox potential of pheophytin a in photosystem II of two cyanobacteria having the different special pair chlorophylls. Proc. Natl. Acad. Sci. U. S. A. 107,3924–3929 (2010).

39. Marquardt, J., Senger, H., Miyashita, H., Miyachi, S. & Mörschel, E. Isolation and characterization of biliprotein aggregates from Acaryochloris marina, a Prochloron-like prokaryote containing mainly chlorophyll d. FEBS Lett. 410,428–432 (1997).

40. Bryant, D. A. et al. Far-red light allophycocyanin subunits play a role in chlorophyll d accumulation in far-red light. Photosynth. Res. 143,81–95 (2020).

41. Gan, F. & Bryant, D.A. Adaptive and acclimative responses of cyanobacteria to far-red light. Environ. Microbiol. 17,3450–3465 (2015).

42. Rippka, R., Deruelles, J., Waterbury, J. B., Herdman, M. & Stanier, R.Y. Generic assignments, strain histories and properties of pure cultures of cyanobacteria. J. Gen. Microbiol. 111,1–61 (1979).

43. Li, Y., Scales, N., Blankenship, R. E., Willows, R. D. & Chen, M. Extinction coefficient for red-shifted chlorophylls: chlorophyll d and chlorophyll f. Biochim. Biophys. Acta - Bioenerg. 1817,1292–1298 (2012).

44. Choubeh, R. R., Wientjes, E., Struik, P. C., Kirilovsky, D. & van Amerongen, H. State transitions in the cyanobacterium Synechococcus elongatus 7942 involve reversible quenching of the photosystem II core. Biochim. Biophys. Acta - Bioenerg. 1859,1059–1066 (2018).

45. Bhatti, A. F., Choubeh, R. R., Kirilovsky, D., Wientjes, E. & van Amerongen, H. State transitions in cyanobacteria studied with picosecond fluorescence at room temperature. Biochim. Biophys. Acta - Bioenerg. 1861,148255 (2020).

46. van Stokkum, I. H. M., van Oort, B., van Mourik, F., Gobets, B. & van Amerongen, H. (Sub)-Picosecond spectral evolution of fluorescence studied with a synchroscan streak-camera system and target analysis. in Biophysical techniques in photosynthesis (eds. Aartsma, T. J. & Matysik, J.) II,223–240 (Springer, 2008).

