## Supplementary Information for "The natural design for harvesting far-red light: the antenna increases both absorption and quantum efficiency of Photosystem II"

### Supplementary Materials

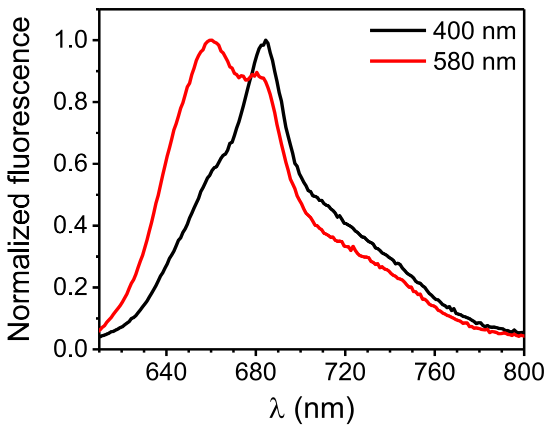


**Figure S1. Fluorescence spectra of WL-cells excited at 400 and 580 nm.** After preferential Chl excitation at 400 nm (black line), steady-state fluorescence shows a peak at ~680 nm due to WL-PSII, whereas WL-PSI is responsible for the shoulder above 700 nm. Upon PBS excitation (580 nm, red line), the emission spectrum shows two partially overlapping peaks at ~660 nm and ~680 nm due to APC and WL-PSII, respectively.

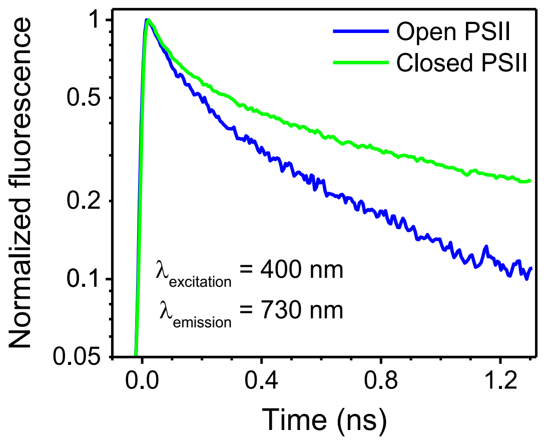


**Figure S2. TRF of FRL-cells.** TRF traces (normalized to their maxima) detected at 730 nm (where both FRL-BCs and FRL-PSII emit) upon 400-nm excitation with mostly open and closed PSII RCs. The presented traces were obtained by integrating the TRF data between 725 and 735 nm and were binned to a time-step of 6 ps. Please note that the fluorescence intensity is on a logarithmic scale.

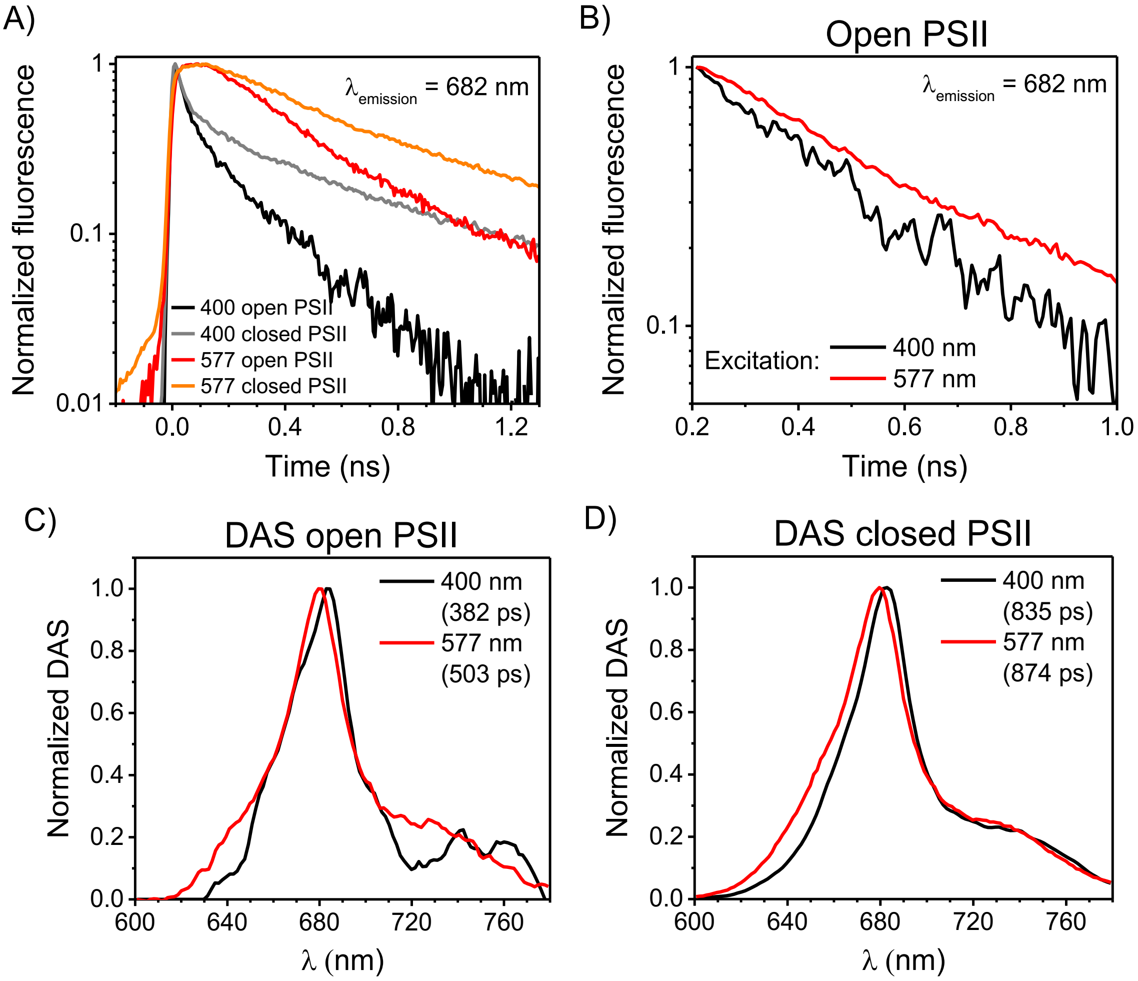


**Figure S3. Time-resolved fluorescence (TRF) of WL-cells.**

A) TRF traces (normalized to their maxima) detected at 682 nm (where WL-PSII emission is maximal) upon 400-nm and 577-nm excitation with mostly open and closed PSII RCs. The presented traces were obtained by integrating the TRF data between 677 and 687 nm and were binned to a time-step of 6 ps. B) TRF traces detected at 682 nm upon 400-nm and 577-nm excitation with mostly open PSII RCs. The traces are normalized to their value around 200 ps, and the first 200 ps of fluorescence kinetics are excluded for clarity. C) Normalized long-lived DAS from TRF measurements with open PSII RCs after 400 and 577 nm excitation (corresponding to the green DAS in Figures S4A and S5A). The differences between the two spectra in the region above 700 nm are mainly due to the higher noise level in the 400-nm excitation dataset, as a consequence of the low powers used in the experiment and the small amplitude of this component. D) Normalized long-lived DAS from TRF measurements with closed PSII RCs after 400-nm and 577-nm excitation (corresponding to the green DAS in Figures S4B and S5B). The lifetimes of the DAS are indicated in parentheses in the legends. The 400-nm excitation data are from Mascoli et al.[1]

The TRF traces of WL-cells detected at 682 nm are substantially longer-lived both after 400-nm and 577-nm excitation when PSII RCs are closed (A). This indicates that a large amount of WL-PBSs (prevalently excited at 577 nm) is energetically connected to WL-PSII. The 682-nm traces with open PSII RCs excited at 400 nm and 577 nm can be used to estimate the kinetics of energy trapping by WL-PSII RCs after either Chl or bilin excitation. A direct comparison of the normalized traces in (A), however, is not informative, as the trace excited at 400 nm also contains a large contribution from the Chls *a* of WL-PSI. However, energy transfer and trapping inside WL-PSI are markedly faster (10-30 ps, as can be seen from the black and red decay associated spectra (DAS) in Figure S4) than excited-state decay in WL-PSII. Furthermore, the average timescale of energy equilibration between the WL-PBS and the photosystems is about 100-150 ps[2–4]. As a result, the fluorescence decay after 200 ps can be assumed to stem mainly from the equilibrated pigments of WL-PSII and the WL-PBSs. When normalized at a delay time of 200 ps, the 682-nm trace excited at 577 nm is longer-lived than that excited at 400 nm (B), indicating a slower trapping of the excitations formed on the bilins than those formed on the Chls. The spectral signatures of WL-PSII with its energetically coupled PBSs can be retrieved from the longest-lived DAS obtained from global analysis of TRF data (see Figures S4-5 for the full set of DAS). This is confirmed by the fact that the lifetime of this DAS substantially increases when PSII RCs close (from 380 ps to 840 ps upon 400-nm excitation, and from 500 ps to 870 ps upon 577-nm excitation). When PSII RCs are open, the lifetime of this DAS also represents an upper estimate of the time required for charge separation to occur. Panel (C) shows that this lifetime rises from less than 400 ps after preferential Chl excitation to about 500 ps after bilin excitation, showing that the diffusion of excitations from the periphery of the WL-PBSs to WL-PSII delays the trapping at the WL-PSII RCs by about 100 ps. On the other hand, the lifetime and spectra of the long-lived DAS excited at 400 nm and 577 nm essentially overlap both in the open and in the closed state (C-D), indicating that excitations in the WL-PBSs and WL-PSII equilibrate to a large extent before decaying.

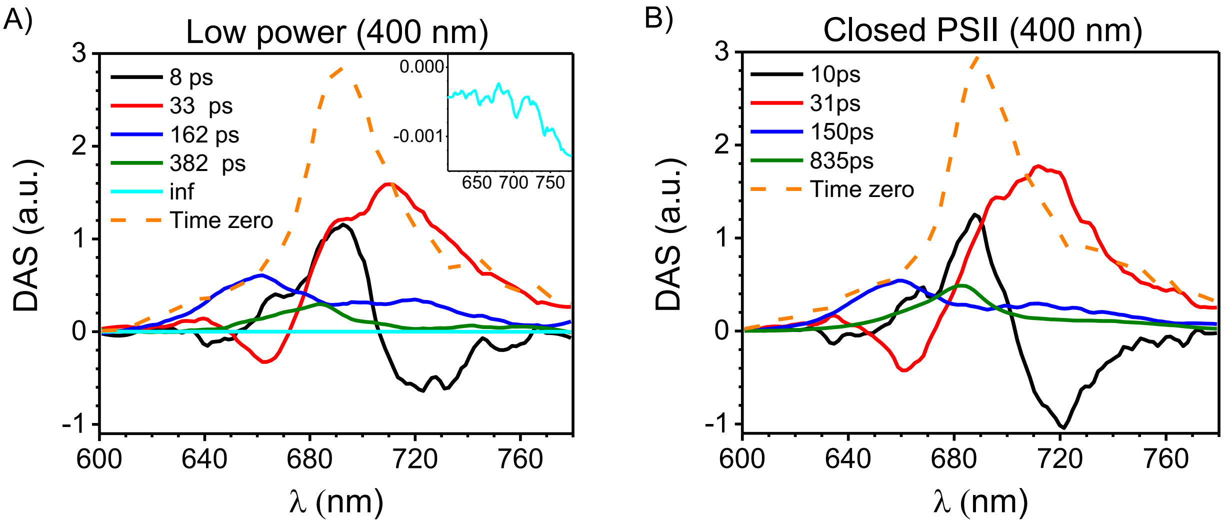


**Figure S4. TRF measurements of WL-cells excited at 400 nm.**

DAS from TRF measurements on WL-cells excited at 400 nm with mostly open (A) and closed (B) PSII RCs (data from Mascoli et al.[1]). The small long-lived (inf) component in (A) (magnified in the inset) reflects some background signal. The time-zero spectrum (orange dashed lines) is calculated as the sum of all other DAS. The DAS are normalized to the area of the time-zero spectrum for better comparison. The two fastest DAS (black and red), with a lifetime of about 10 and 30 ps, respectively, can be largely ascribed to downhill energy transfer and trapping by WL-PSI. At shorter wavelengths, smaller band shift features due to downhill energy transfer in the WL-PBS are also observed (as bilins are excited to a minor extent at this wavelength). The ~150 ps DAS (blue) is entirely positive and stems prevalently from the decay of bilin excitations The amplitude of this DAS at 680 nm slightly decreases in favor of the longer-lived component when PSII RCs close, indicating some contribution from WL-PSII. The long-lived component (green), whose lifetime and amplitude increase in closed state, can be attributed to WL-PSII, with some WL-PBS contribution at shorter wavelengths. The ratio between the area of the long-lived PSII DAS and that of the time-zero spectrum in closed state represents a low estimate for the amount of initial excitations that are emitted by WL-PSII (as the shorter-lived DAS also contain minor contributions from WL-PSII). The amplitude of the longest-lived PSII DAS is much smaller than that of the 30 ps DAS (representing WL-PSI decay) and accounts for only 15% of the total initial excitations (based on the area of the time-zero spectrum, orange dashed lines). This implies that the largest amount of initial Chl excitations is trapped by the RCs of PSI and is in agreement with the fact that PSI commonly binds the majority of total Chls in cyanobacterial cells[5,6].

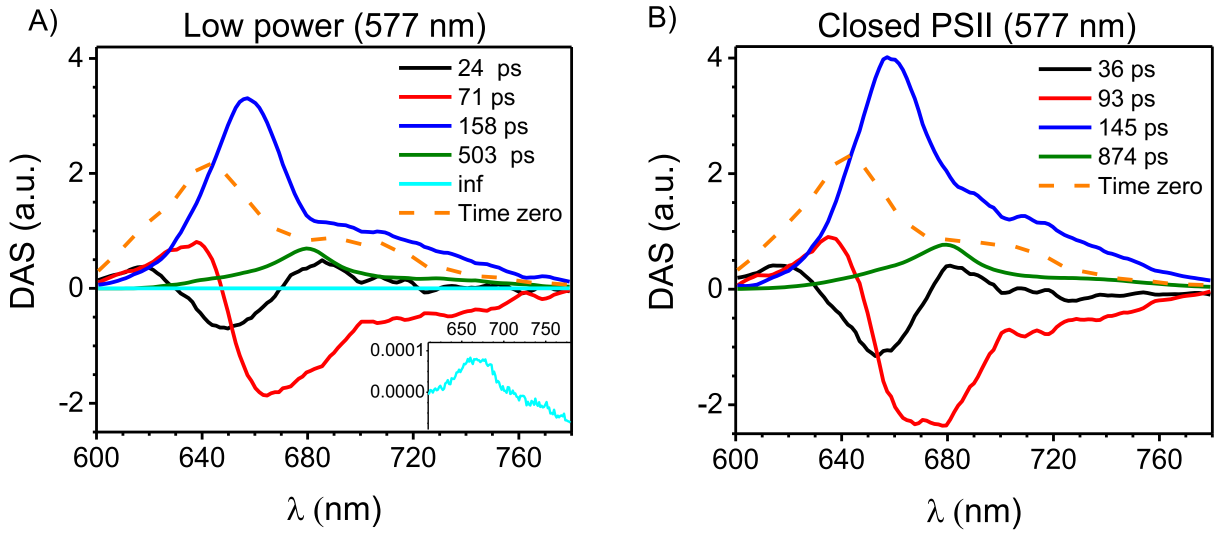


**Figure S5. TRF measurements of WL-cells excited at 577 nm.**

A,B) DAS from TRF measurements on WL-cells excited at 577 nm with mostly open (A) and closed (B) PSII RCs. The small long-lived (inf) component in (A) (magnified in the inset) reflects some background signal and, possibly, a vanishingly small amount of unconnected PBS/closed PSII. The time-zero spectrum (orange dashed lines) is calculated as the sum of all other DAS. The DAS are normalized to the area of the time-zero spectrum for better comparison. The first three DAS (black, red and blue), whose lifetime and shape are very similar in open and closed state, stem mostly from the phycobiliproteins. The first two DAS describe the funneling of excitations from the peripheral rods to the inner core of WL-PBSs, occurring in less than 100 ps, and might also include energy transfer from the PBS core to the photosystems. The third DAS (blue), with a lifetime of about 150 ps, is entirely positive and represents, therefore, an excited-state decay. It is also very similar to the ~150 ps DAS observed upon 400 nm excitation (Figure S4, blue), suggesting a common origin. This DAS can be largely ascribed to WL-PBS decay, although contributions from the two photosystems are probably present at $\lambda \geq 680$ nm. This decay is substantially shorter than in isolated PBSs (typically ns)[7] due to the connectivity of the PBS to PSI and PSII (photochemical quenching)[8,9] *in vivo*. The lifetime of ~150 ps, which is comparable to the typical migration time from the PBS to the photosystem cores[2–4], and the prevalent PBS contribution to these DAS, support the hypothesis that trapping of the PBS excitations by the RCs is to a large extent migration limited. This is particularly true for the WL-PBS connected to WL-PSI, as the latter can perform charge separation on a faster timescale than that of energy migration (~30 ps, see Figure S4). Furthermore, the weak dependency on the state of PSII RCs implies that the ~150 ps DAS largely stems from WL-PBSs that transfer excitations to WL-PSI. At the same time, some contribution from WL-PBSs quenched by WL-PSII RCs might also be present, since the fluorescence decay of PSII in cyanobacteria is inherently multi-exponential and displays relatively short-lived components also in the closed state[10]. The slowest DAS (green), whose lifetime increases when PSII RCs close, can be attributed to WL-PSII connected to WL-PBSs. The ratio between the area of the green DAS and that of the time-zero spectrum is 0.29. When accounting for the fact that the oscillator strength of the Chls in WL-PSII is only about 60% of that of the initially excited bilins[11,12], it follows that at least 50% of the initial WL-PBS excitations (577 nm) are transferred to WL-PSII, which is similar to what found in *Synechocystis*[8]. This value is much larger than the 15% of initial excitations emitted by WL-PSII after Chl excitation (400 nm). This finding confirms that PBSs increase the antenna size of PSII relative to PSI[2,13]. The results presented so far for WL-cells are also consistent with those obtained previously for *Synechocystis* sp. PCC6803[14,15] and *Synechococcus elongatus*[9].

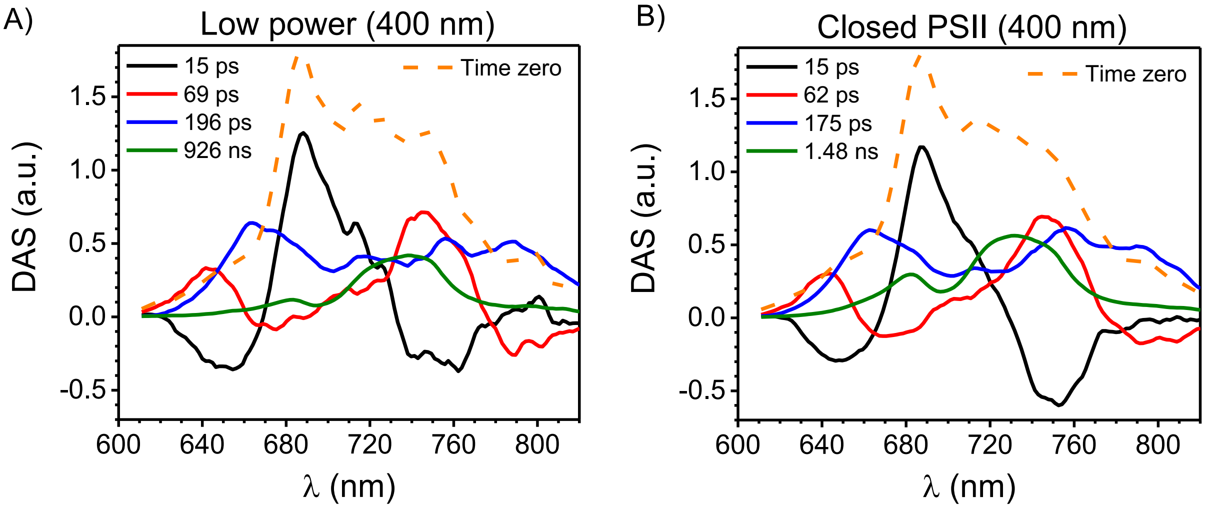


**Figure S6. TRF measurements of FRL-cells excited at 400 nm.**

DAS from TRF measurements on FRL-cells excited at 400 nm with mostly open (A) and closed (B) PSII RCs. The time-zero spectrum (orange dashed lines) is calculated as the sum of all other DAS. The DAS are normalized to the area of the time-zero spectrum for better comparison. The first two components mostly describe downhill energy transfer from Chl *a* to Chl *f* (in both FRL-PSI and FRL-PSII, black DAS) and from the majority of Chls *f* to strongly red-shifted Chls *f* (in FRL-PSI only, red DAS)[1], respectively. The 400-nm laser also excites the bilins to some extent, as witnessed by the downhill energy transfer features observed in both DAS at shorter wavelengths (similar features dominate the early kinetics when exciting the phycobiliproteins prevalently, see Figure S7). A third component, with a lifetime < 200 ps (blue lines), can be largely assigned to FRL-PSI trapping (in the 750-800 nm region) and WL-PBS/WL-PSII decay (in the 640-690 nm region). Some contribution from FRL-BCs and FRL-PSII is presumably present at intermediate wavelengths (710-740 nm). The overall shape/amplitude of this component is not very sensitive to the state of PSII RCs, though the relative amplitude around 680 nm decreases in closed state in favor of the longer-lived components (*cfr* blue and green DAS in (A) and (B)), confirming the contribution from WL-PSII at these wavelengths. The long-lived DAS (green) increases in amplitude and lifetime (from less than 1 ns in the open state to almost 1.5 ns in the closed state) when PSII RCs close and can be mostly ascribed to WL-PSII (with a peak at 680 nm) and FRL-PSII (with a peak at 735-740 nm). As we showed in our previous work[1], the spectrum of FRL-PSII *in vivo* has more amplitude at shorter wavelengths in comparison to isolated FRL-PSII due to the contribution of FRL-BCs, advocating for an association between these two units in the cells. This substantial emission by FRL-BCs after 400-nm excitation might result from their direct excitation and, possibly, from a partial equilibration of the excitations formed in FRL-PSII with the red-shifted APC of FRL-BCs. Energy transfer from FRL-PSII to FRL-BCs is probably more favored than that from WL-PSII to the WL-PBS core, for two reasons: (i) an entropic factor due to the small number of red-shifted Chls in FRL-PSII and (ii) a kinetic factor, as the excitations in FRL-PSII power photochemistry more slowly than in WL-PSII and have more time to equilibrate with the nearby APC. The ratio between the area of the long-lived PSII-related DAS in closed state (green line in (B)) and that of the time zero spectrum (orange dashed line) is about 0.28 for FRL-cells, which is almost two times that observed in WL-cells (0.15). This suggests that FRL-cells have more PSII relative to PSI in comparison to WL-cells. Note that the data presented here are very similar to those in our previous work[1]. It is also likely that, in our measurement at low power, a small amount of WL-PSII with closed RCs is present (as the lifetime of the longest-lived component, i.e. ~930 ps, is similar to the one observed for WL-PSII with closed RCs in the WL-cells (Figure S5B). On the other hand, FRL-PSII RCs are fully open in the measurement at low power, as confirmed by the data on power dependency shown in Figure S11.

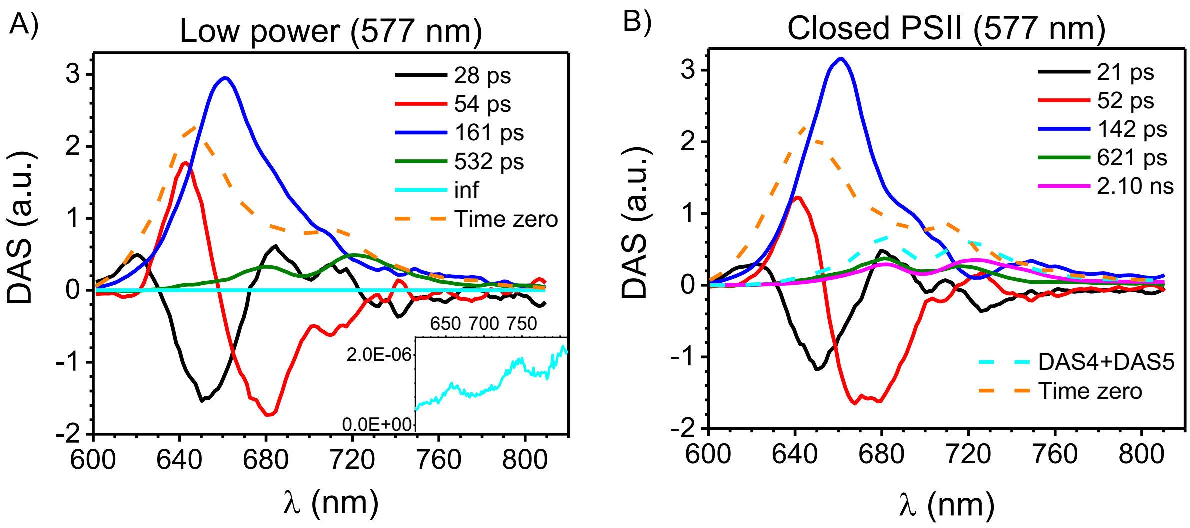


**Figure S7. TRF measurements of FRL-cells excited at 577 nm.**

DAS from TRF measurements on FRL-cells excited at 577 nm with mostly open (A) and closed (B) PSII RCs. The small long-lived (inf) component in (A) (magnified in the inset) reflects some background signal. The time-zero spectrum (orange dashed lines) is calculated as the sum of all other DAS. The DAS are normalized to the area of the time-zero spectrum for better comparison. According to the steady-state fluorescence spectra, FRL-cells contain WL-PBSs and FRL-BCs, which are both excited at 577 nm (with a preference for phycocyanin and, therefore, for the WL-PBSs). Note that it is extremely difficult to excite FRL-BCs with a higher selectivity, as their absorption maxima (around 650 nm and 710 nm)[16,17] overlap with the absorption bands of the Chls (and with the detection window) to a much larger extent. Similar to what was observed for WL-cells (Figure S5), the first two DAS (black and red lines) are dominated by downhill energy transfer in the WL-PBS and from them, possibly, to WL-PSII. Some energy transfer to far-red emitting units (FRL-BCs and FRL-photosystems) might also contribute to these components. The third DAS (blue lines), with a lifetime of ~150 ps, is weakly sensitive to the state of the PSII RCs and reminiscent of the ~150 ps DAS observed in WL-cells (blue lines in Figure S5). This component largely represents photochemical quenching of WL-PBS excitations by the RCs of WL-photosystems (trapping by FRL-photosystems can be excluded, as it is substantially slower)[1]. Its amplitude relative to the time-zero spectrum, however, is reduced in FRL-cells (as the 577-nm laser also excites FRL-BCs, which decay more slowly and contribute to the longer-lived DAS, see below). The ~150 ps DAS in FRL-cells also have a lower relative amplitude above 700 nm and show a dip at about 725 nm (more evident in closed state, (B)), indicating that some energy transfer to the FRL-units also takes place on a similar time scale. Some contribution from WL-PSII is probably present at 680 nm, as the DAS amplitude at this wavelength decreases in favor of the longer-lived components when PSII RCs close (*cfr* blue and green/magenta lines in open/closed state). The long-lived components are mostly associated with fluorescence from WL-PSII (peaking at 680 nm) connected to WL-PBSs and, at longer wavelengths, FRL-BCs and FRL-PSII. In open state (A), the lifetime of the long-lived DAS (green) is ~530 ps, about 400 ps shorter than the longest-lived DAS observed after 400-nm excitation (Figure S6A). This implies that trapping of bilin excitations by FRL-PSII RCs is faster than trapping of Chl excitations. Notably, the lifetimes of these two components (~530 ps after bilin excitation, and ~930 ps upon Chl excitation) are estimated from the global analysis across the entire spectral range, and are therefore influenced by the timescales of WL-PSII decay (as the system is highly heterogeneous). To eliminate the contribution of WL-units from the estimation of the longest lifetimes, we also performed global analysis on restricted datasets including only wavelengths > 700 nm. Figure S8 shows the results of these analyses which, despite some minor differences in the fitted lifetimes, are fully consistent with our previous conclusions (i.e. trapping by FRL-PSII RCs after bilin excitation is faster than after Chl excitation).

In the closed state (B), two long-lived DAS can be resolved: one has a lifetime of about 600 ps (green) and is enriched in WL-PSII and FRL-BCs, the other (magenta), with a lifetime above 2 ns, contains contributions from WL-PSII, FRL-BCs and FRL-PSII. In agreement with the substantial variable fluorescence shown by the 730-nm emission traces after 577-nm excitation (Figure 2A), the average lifetime of the long-lived DAS also increases when PSII RCs close. These DAS have a large contribution from red-shifted APC, implying that FRL-BCs are largely connected to FRL-PSII. Note that, in principle, FRL-BCs might also be connected to WL-PSII. However, though this possibility cannot be completely excluded, it seems less likely because the presence of a large uphill energy barrier between red-shifted APC and Chl *a* is expected to slow down trapping in the RCs of WL-PSII to a large extent. However, the longest-lived decay of FRL-BCs in open state is not much longer than that of PSII of WL-cells. The amount of phycobiliproteins connected to FRL-PSI is also likely to be small since all 577-nm excited DAS have a very low amplitude above 750 nm. Overall, the observation that WL-PBSs remain preferentially connected to WL-photosystems, whereas FRL-BCs are coupled to FRL-photosystems (prevalently FRL-PSII) is in agreement with the spatial segregation between FRL- and WL-units observed in another FaRLiP strain[18].

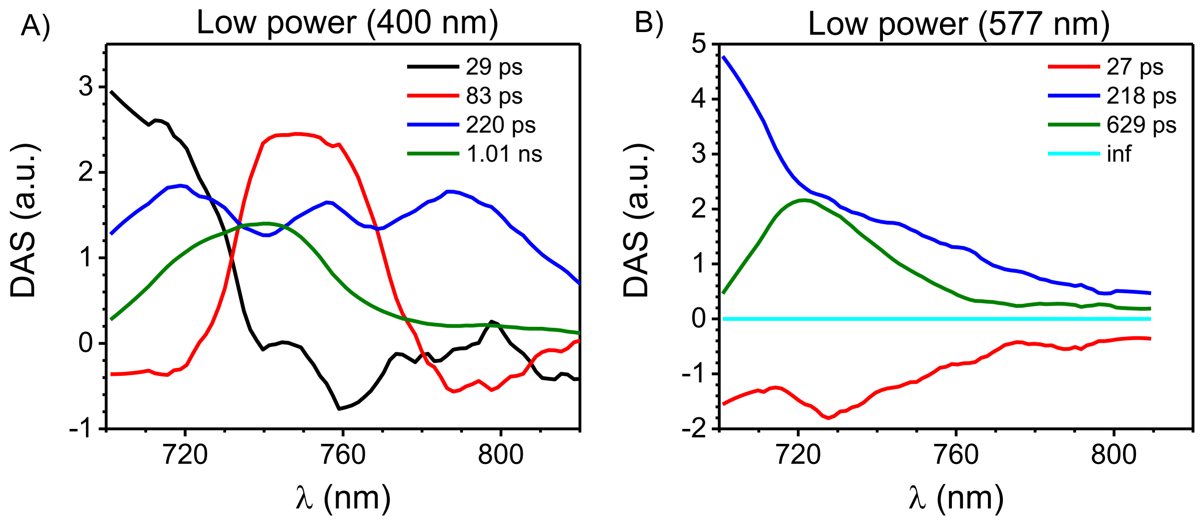


**Figure S8. TRF measurements of FRL-cells with open PSII-RCs.**

DAS from TRF measurements on FRL-cells excited at 400 nm (A) and 577 nm (B) with mostly open PSII RCs. The global analysis has been restricted to the spectral region above 700 nm to minimize the influence of the kinetics of WL-units from the estimation of the excited-state decay lifetimes of FRL-units.

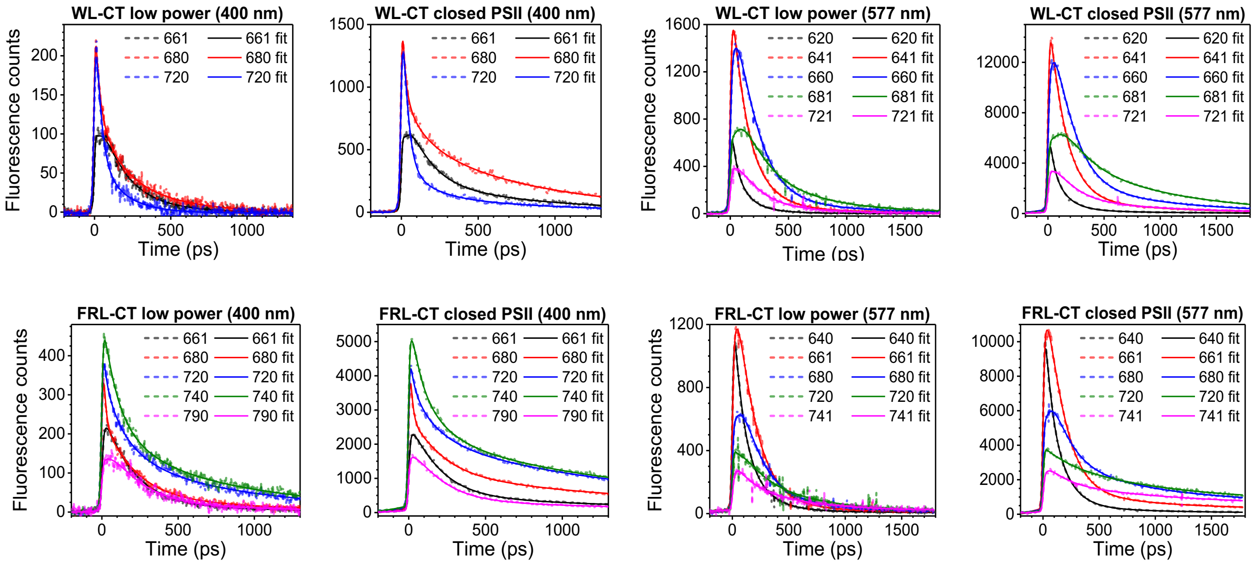


**Figure S9. TRF measurements: raw data versus fittings.**

First row: overlay of raw and globally fitted fluorescence time traces for WL-cells upon 400-nm and 577-nm excitation at selected emission wavelengths (see Figures S4-5 for the DAS from global analysis). The selected emission wavelengths highlight contributions from short-wavelength phycocyanin (620 nm), long-wavelength phycocyanin (641 nm), APC (660 nm), WL-PSII (680-681 nm), and WL-PSI (720-721 nm).

Second row: overlay of raw and globally fitted fluorescence time traces for FRL-cells upon 400-nm and 577-nm excitation at selected emission wavelengths (see Figures S6-7 for the DAS from global analysis). The selected emission wavelengths highlight contributions from phycocyanin (640 nm), APC (661 nm), WL-PSII (680 nm), red-shifted APC (720 nm), FRL-PSII (740-741 nm), and FRL-PSI (790 nm).

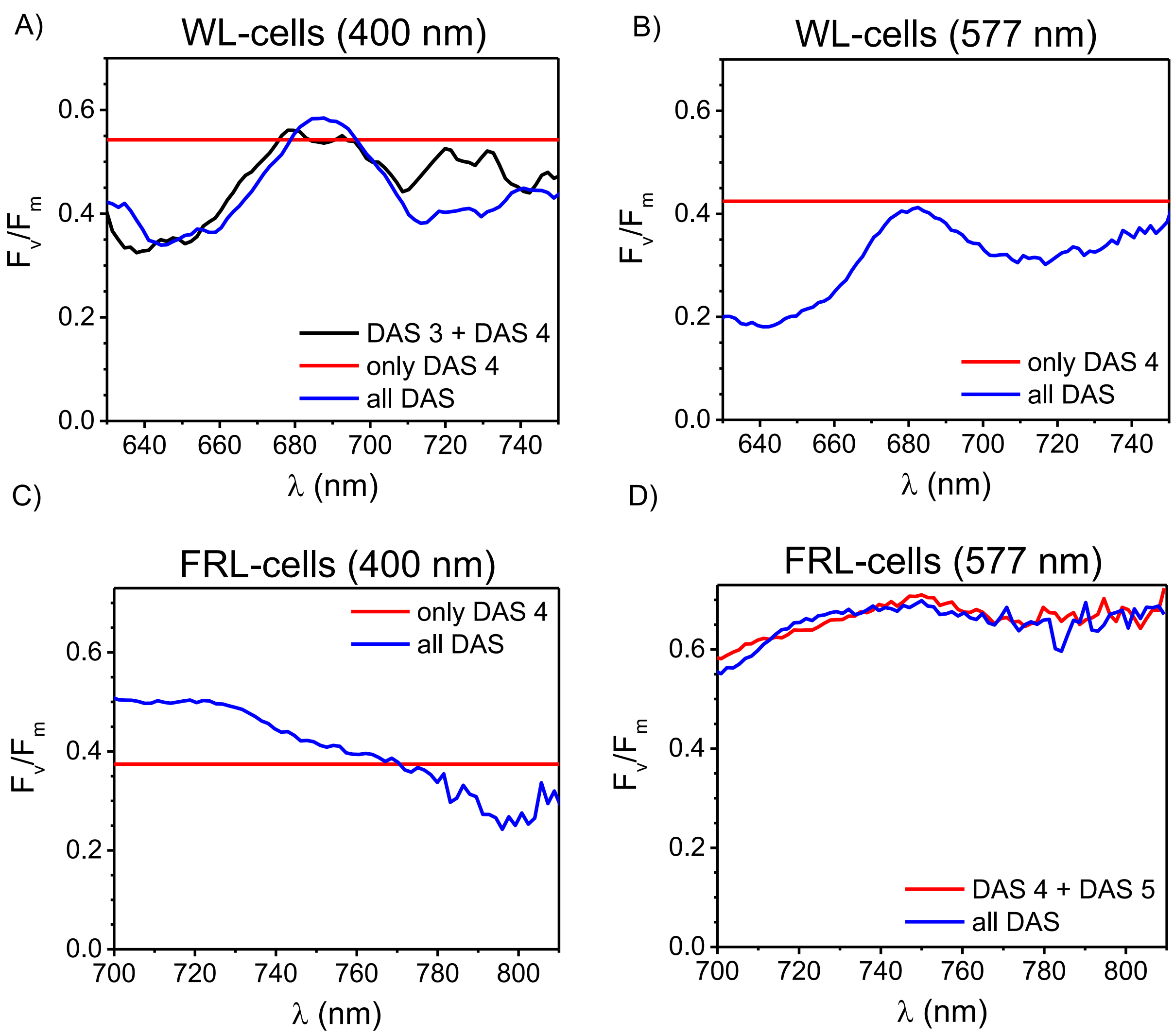


**Figure S10. PSII photochemical efficiency.**

$F_v/F_m$ of WL-cells at different wavelengths calculated with different methods from the DAS of TRF measurements (Figures S4-7). The aim of these calculations is to evaluate the impact of PSI fluorescence on the estimation of the PSII efficiency. For this purpose, we compared the $F_v/F_m$ obtained from the whole TRF data (i.e. using all the DAS, as in Figure 3) to that obtained based on the longest-lived components only, as these components are mostly contributed by PSII (and, possibly, the phycobiliproteins). These two levels of approximation represent opposite extremes, as in the first case the contribution of PSI is also included in the estimation of the variable fluorescence (which is, in principle, a property of PSII only), whereas in the latter case, the contribution of PSII to the shortest-lived DAS (which is generally not zero) is neglected. In the following, the average lifetime is obtained at each wavelength from the DAS as: $\tau_{avg}(\lambda) = (\sum_k DAS_k(\lambda) \cdot \tau_k) / \sum_k DAS_k(\lambda)$, where *k* designates the DAS that participate in the average.

A,B) WL-cells upon different excitations: in (A), the black line shows $F_v/F_m$ obtained as $(1 - \tau_o/\tau_m)$, where $\tau_o$ and $\tau_m$ are the average lifetimes in open and closed state, respectively, calculated using only the DAS 3 and 4 of Figure S4 (those with the longest lifetime), which do not contain contributions from PSI (both in the open and in the closed state). The red line displays $1 - \tau_o/\tau_m$, where only the longest lifetime has been used, for both the open and the closed state (as the longest-lived DAS in Figure S4 mostly contain contributions from PSII). The blue line is obtained by using all DAS from both measurements in Figure S4 (identical to what shown in Figure 3C). The data in (B) are obtained in a similar way from the DAS in Figure S5, using only the longest-lived DAS (red line), which are mostly contributed by PSII, and all DAS (blue lines), as also shown in Figure 3C of the manuscript.

C,D) $F_v/F_m$ of FRL-cells at different wavelengths calculated with different methods from the DAS of TRF measurements (Figures S6-7). In (C), the red line is obtained as in the previous plots, using only the lifetimes of the longest-lived DAS of Figure S6 in open and closed state (which are mostly contributed by FRL-PSII; note that FRL-PSII also contributes to the shorter-lived DAS to some extent), while the blue line is obtained using all DAS (as in Figure 3F). In (D), the red line is obtained from the average lifetimes in open and closed state calculated from the longest-lived DAS in open state (Figure S7A) and the two longest-lived DAS in closed state (Figure S7B). The blue line shows $F_v/F_m$ calculated from all DAS as in Figure 3F.

These plots show that, even though the precise $F_v/F_m$ values can depend on the level of approximation, the qualitative trends are always the same: i.e. $F_v/F_m$ of bilin excitations is lower than that of Chl excitations in the PSII of WL-cells (compare lines of corresponding colors in (A) and (B)), while $F_v/F_m$ of bilin excitations is higher than that of Chl excitations in the FRL-PSII (compare lines of corresponding colors in (C) and (D)).

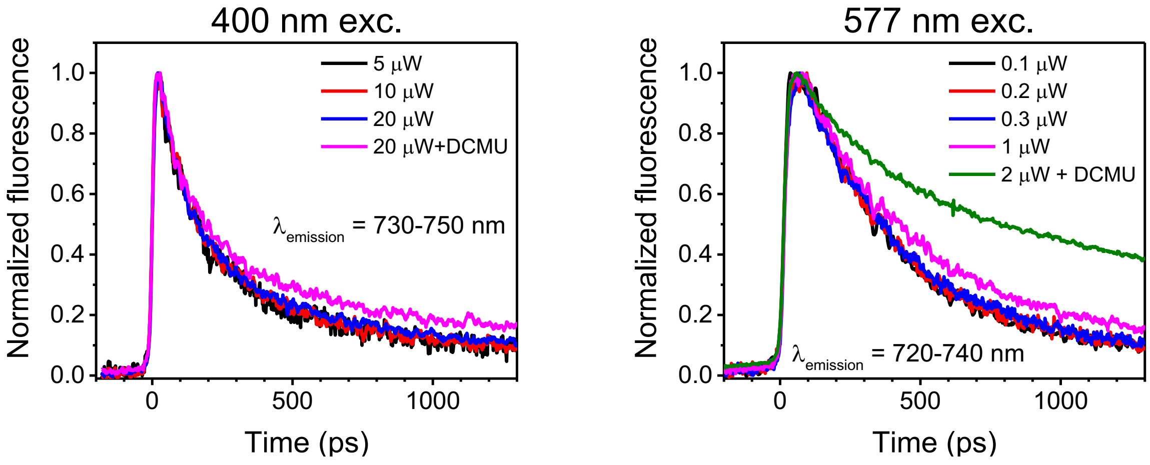


**Figure S11. Power dependency of TRF measurements on FRL-cells.** TRF traces of FRL-cells recorded between 730-750 nm (maximal FRL-PSII emission upon 400-nm excitation) and 720-740 nm (maximal FRL-BC/FRL-PSII emission upon 577-nm excitation) at different excitation powers. The TRF traces are nearly power independent below 10 μW (upon 400-nm excitation) and 0.3 μW (upon 577-nm excitation) and become increasingly longer-lived at higher powers due to higher amounts of closed PSII RCs. In order to keep FRL-PSII RCs open, measurements were performed at 5 μW and 0.2 μW for 400-nm and 577-nm excitation experiments, respectively.

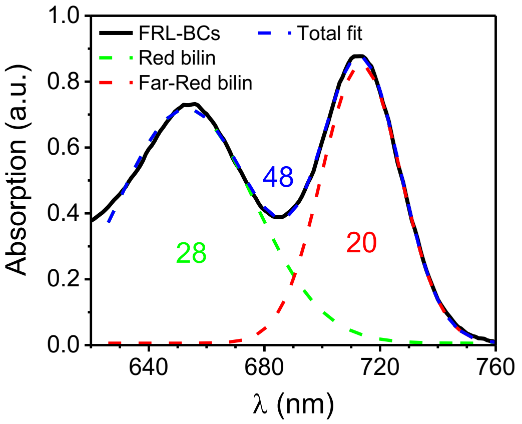


**Figure S12. Estimation of the number of FRL-absorbing bilins in FRL-BCs.** Gaussian fitting of the absorption spectrum of purified FRL-BCs of the FaRLiP strain *Halomicronema hongdechloris* from Li et al.[17]. The fitting is restricted to the spectral region above 630 nm and returns two main Gaussian components: a broader one peaking at 653 nm and ascribed to red-light-absorbing bilins, and a narrower one peaking at 713 nm and ascribed to FRL-absorbing bilins. The areas under the two Gaussians can be used to estimate the size of both pigment pools. When normalizing to a total of 48 bilins (24 per each core cylinder)[2,16], it is found that FRL-BCs contain approximately 20 FRL-absorbing bilins. Note that the absorption spectrum of FRL-BCs from another FaRLiP strain, *Synechococcus* sp. PCC 7335[16], is very similar to the one from *Halomicronema hongdechloris* shown here, implying that the number of FRL-absorbing bilins in FRL-BCs does not change much in different species. Furthermore, when calculating the antenna size under FRL, only the FRL-absorbing pigments are taken into account, meaning that our calculations in the main text are not influenced by the precise organization of phycobiliproteins under FRL (i.e. whether the FRL-BCs attach to other phycobiliprotein units or not).

| Subunit | Chl | Coupling with $Chl_{D1}$ ($cm^{-1}$) | Hopping time to $Chl_{D1}$ (ps) | Coupling with $P_{D2}$ ($cm^{-1}$) | Hopping time to $P_{D2}$ (ps) |
|---|---|---|---|---|---|
| D1 | $P_{D1}$ | -18.2 | 3.9.E+00 | 181.6 | 3.9.E-02 |
| D1 | $Chl_{D1}$ | - | | -90.9 | 1.6.E-01 |
| D1 | $Chl_{D2}$ | 14.1 | 6.5.E+00 | -9.6 | 1.4.E+01 |
| D1 | $Chl_{Z1}$ | 2.9 | 1.5.E+02 | 1.3 | 7.9.E+02 |
| D2 | $P_{D2}$ | -90.9 | 1.6.E-01 | - | |
| D2 | $Chl_{Z2}$ | -0.1 | 4.1.E+05 | 1.0 | 1.3.E+03 |
| CP47 | 602 | -0.1 | 1.2.E+05 | 0.1 | 1.3.E+05 |
| CP47 | 603 | 1.1 | 1.1.E+03 | 1.5 | 5.7.E+02 |
| CP47 | 604 | -0.4 | 9.7.E+03 | -0.9 | 1.8.E+03 |
| CP47 | 605 | -0.6 | 4.2.E+03 | 1.6 | 5.0.E+02 |
| CP47 | 606 | 1.3 | 7.4.E+02 | 1.1 | 9.8.E+02 |
| CP47 | 607 | -0.8 | 2.0.E+03 | -0.9 | 1.5.E+03 |
| CP47 | 608 | -2.3 | 2.5.E+02 | **-3.8** | **8.8.E+01** |
| CP47 | 609 | -2.3 | 2.4.E+02 | -2.1 | 2.8.E+02 |
| CP47 | 610 | -0.9 | 1.8.E+03 | -1.1 | 1.0.E+03 |
| CP47 | 611 | -1.1 | 1.2.E+03 | -1.7 | 4.7.E+02 |
| CP47 | 612 | 0.6 | 3.2.E+03 | -3.0 | 1.5.E+02 |
| CP47 | 613 | 0.2 | 3.2.E+04 | 1.1 | 1.0.E+03 |
| CP47 | 614 | -1.3 | 7.2.E+02 | -2.9 | 1.5.E+02 |
| CP47 | 615 | -1.4 | 6.2.E+02 | -2.2 | 2.6.E+02 |
| CP47 | 616 | 0.3 | 1.3.E+04 | -0.2 | 3.7.E+04 |
| CP47 | 617 | -0.6 | 3.3.E+03 | -0.6 | 3.7.E+03 |
| CP43 | 501 | 2.0 | 3.3.E+02 | 1.1 | 1.1.E+03 |
| CP43 | 502 | -2.4 | 2.3.E+02 | -0.4 | 8.5.E+03 |
| CP43 | 503 | -0.3 | 2.0.E+04 | -0.9 | 1.5.E+03 |
| CP43 | 504 | 1.8 | 3.9.E+02 | 2.6 | 2.0.E+02 |
| CP43 | 505 | **7.5** | **2.3.E+01** | **4.0** | **7.9.E+01** |
| CP43 | 506 | 0.5 | 4.7.E+03 | 0.8 | 2.0.E+03 |
| CP43 | 507 | 2.6 | 1.8.E+02 | 1.4 | 6.2.E+02 |
| CP43 | 508 | -0.8 | 2.2.E+03 | -2.3 | 2.3.E+02 |
| CP43 | 509 | -1.6 | 5.2.E+02 | -0.3 | 1.3.E+04 |
| CP43 | 510 | 2.1 | 2.8.E+02 | 1.1 | 9.8.E+02 |
| CP43 | 511 | 0.3 | 1.7.E+04 | 0.1 | 3.8.E+05 |
| CP43 | 512 | -0.3 | 1.1.E+04 | -0.8 | 2.2.E+03 |
| CP43 | 513 | 0.8 | 2.2.E+03 | 0.7 | 2.6.E+03 |

**Table S1. Electronic couplings and excitation energy transfer hopping times between all Chls in the PSII core and the primary electron donor in the RC (either $Chl_{D1}$ or $P_{D2}$).** The couplings between Chl $Q_y$ transitions are based on the 1.9Å-resolution crystal structure of PSII core of *Thermosynechococcus vulcanus* (PDB: 3WU2)[19] and were calculated on a single PSII monomer using the point dipole approximation as in Mascoli et al.[1]. Both the excitation energy transfer (EET) donor and acceptor where assumed to be Chls *f* and their oscillator strength was set to be 1.5 times that of Chl *a*[20]. Hopping times were calculated in the Förster approximation, assuming that the EET donor in the antenna emits at 740 nm (based on fluorescence spectra of FRL-PSII)[1], while the EET acceptor (i.e. the primary electron donor in the RC) absorbs at 727 nm[21,22]. The spectral shapes were assumed to be Gaussians with σ = 100 $cm^{-1}$, which is reasonable at RT. The entries with couplings and hopping times involving the Chls in the PSII core antenna (CP43/47) as EET donors are highlighted in green (when the EET acceptor is $Chl_{D1}$) and yellow (when the EET acceptor is $P_{D2}$). The Chls in the antenna (CP43/47) resulting in hopping times shorter than 100 ps are highlighted in bold. When the primary electron donor is $Chl_{D1}$ (which is spatially closer to CP43), the Chls of CP43 are generally better connected to the RC than those of CP47 (see also Figure 4). Furthermore, Chl 505, facing the stroma, displays by far the fastest EET to the RC. On the other hand, when the primary electron donor is $P_{D2}$, there is no large difference in connectivity (on average) between CP43 or CP47 and the RC. Additionally, Chl 505 of CP43 and Chl 608 of CP47 both have hopping times shorter than 100 ps. Chl 608, however, is closer to the lumen and is, therefore, not a good candidate for bridging the RC of FRL-PSII to the FRL-BCs, which should attach to PSII on the stromal side. This leaves Chl 505 as the best option for connecting FRL-BCs and FRL-PSII RCs even when $P_{D2}$ functions as the primary electron donor.

#### References


1. Mascoli, V., Bersanini, L. & Croce, R. Far-red absorption and light-use efficiency trade-offs in chlorophyll f photosynthesis. *Nat. Plants* **6,** 1044–1053 (2020).
2. Bryant, D. A. & Canniffe, D. P. How nature designs light-harvesting antenna systems: design principles and functional realization in chlorophototrophic prokaryotes. *J. Phys. B At. Mol. Opt. Phys.* **51,** 33001 (2018).
3. van Grondelle, R., Dekker, J. P., Gillbro, T. & Sundström, V. Energy transfer and trapping in photosynthesis. *Biochim. Biophys. Acta* **1187,** 1–65 (1994).
4. Akhtar, P. *et al.* Time-resolved fluorescence study of excitation energy transfer in the cyanobacterium Anabaena PCC 7120. *Photosynth. Res.* **144,** 247–259 (2020).
5. Manodori, A. & Melis, A. Cyanobacterial acclimation to photosystem I or photosystem II light. *Plant Physiol.* **82,** 185–189 (1986).
6. Fujita, Y. A study on the dynamics features of photosystem stoichiometry: accomplishments and problems for future studies. *Photosynth. Res.* **53,** 83–93 (1997).
7. Tian, L. *et al.* Picosecond kinetics of light harvesting and photoprotective quenching in wild-type and mutant phycobilisomes isolated from the cyanobacterium Synechocystis PCC 6803. *Biophys. J.* **102,** 1692–1700 (2012).
8. Tian, L. *et al.* Site, rate, and mechanism of photoprotective quenching in cyanobacteria. *J. Am. Chem. Soc.* **133,** 18304–18311 (2011).
9. Bhatti, A. F., Choubeh, R. R., Kirilovsky, D., Wientjes, E. & van Amerongen, H. State transitions in cyanobacteria studied with picosecond fluorescence at room temperature. *Biochim. Biophys. Acta - Bioenerg.* **1861,** 148255 (2020).
10. Tian, L., Farooq, S. & van Amerongen, H. Probing the picosecond kinetics of the photosystem II core complex in vivo. *Phys. Chem. Chem. Phys.* **15,** 3146–3154 (2013).
11. Cohen-Bazire, G., Béguin, S., Rimon, S., Glazer, A. N. & Brown, D. M. Physico-chemical and immunological properties of allophycocyanins. *Arch. Microbiol.* **111,** 225–238 (1977).
12. Porra, R. J., Thompson, W. A. & Kriedemann, P. E. Determination of accurate extinction coefficients and simultaneous equations for assaying chlorophylls a and b extracted with four different solvents: verification of the concentration of chlorophyll standards by atomic

absorption spectroscopy. *Biochim. Biophys. Acta* **975,** 384–394 (1989).

13. Glazer, A. N. Phycobilisome. A macromolecular complex optimized for light energy transfer. *Biochim. Biophys. Acta* **768,** 29–51 (1984).

14. Krumova, S. B. *et al.* Monitoring photosynthesis in individual cells of synechocystis sp. PCC 6803 on a picosecond timescale. *Biophys. J.* **99,** 2006–2015 (2010).

15. Chukhutsina, V., Bersanini, L., Aro, E. M. & van Amerongen, H. Cyanobacterial flv4-2 operon-encoded proteins optimize light harvesting and charge separation in photosystem II. *Mol. Plant* **8,** 747–761 (2015).

16. Ho, M.-Y., Gan, F., Shen, G. & Bryant, D. A. Far-red light photoacclimation (FaRLiP) in Synechococcus sp. PCC 7335. II. Characterization of phycobiliproteins produced during acclimation to far-red light. *Photosynth. Res.* **131,** 187–202 (2017).

17. Li, Y. *et al.* Characterization of red-shifted phycobilisomes isolated from the chlorophyll f-containing cyanobacterium Halomicronema hongdechloris. *Biochim. Biophys. Acta - Bioenerg.* **1857,** 107–114 (2016).

18. Majumder, E. L. W. *et al.* Subcellular pigment distribution is altered under far-red light acclimation in cyanobacteria that contain chlorophyll f. *Photosynth. Res.* **134,** 183–192 (2017).

19. Umena, Y., Kawakami, K., Shen, J. R. & Kamiya, N. Crystal structure of oxygen-evolving photosystem II at a resolution of 1.9 Å. *Nature* **473,** 55–60 (2011).

20. Li, Y., Scales, N., Blankenship, R. E., Willows, R. D. & Chen, M. Extinction coefficient for red-shifted chlorophylls: chlorophyll d and chlorophyll f. *Biochim. Biophys. Acta - Bioenerg.* **1817,** 1292–1298 (2012).

21. Nürnberg, D. J. *et al.* Photochemistry beyond the red limit in chlorophyll f-containing photosystems. *Science* **360,** 1210–1213 (2018).

22. Judd, M. *et al.* The primary donor of far-red photosystem II: ChlD1 or PD2? *Biochim. Biophys. Acta - Bioenerg.* **1861,** 148248 (2020).
